# Evolving neoantigen profiles in colorectal cancers with DNA repair defects

**DOI:** 10.1101/667097

**Authors:** Giuseppe Rospo, Annalisa Lorenzato, Nabil Amirouchene-Angelozzi, Alessandro Magrì, Carlotta Cancelliere, Giorgio Corti, Carola Negrino, Vito Amodio, Monica Montone, Alice Bartolini, Ludovic Barault, Luca Novara, Claudio Isella, Enzo Medico, Andrea Bertotti, Livio Trusolino, Giovanni Germano, Federica Di Nicolantonio, Alberto Bardelli

## Abstract

**Background:** Neoantigens that arise as a consequence of tumour-specific mutations can be recognized by T lymphocytes leading to effective immune surveillance. In colorectal cancer (CRC) and other tumour types, a high number of neoantigens is associated with patient response to immune therapies. The molecular processes governing the generation of neoantigens and their turnover in cancer cells are poorly understood. We exploited CRC as a model system to understand how alterations in DNA repair pathways modulate neoantigen profiles over time.

**Methods:** We performed Whole Exome Sequencing (WES) and RNA sequencing (RNAseq) in CRC cell lines, *in vitro* and *vivo*, and in CRC patient-derived xenografts (PDXs) to track longitudinally genomic profiles, clonal evolution, mutational signatures and predicted neoantigens.

**Results:** The majority of CRC models showed remarkably stable mutational and neoantigen profiles, however those carrying defects in DNA repair genes continuously diversified. Rapidly evolving and evolutionary stable CRCs displayed characteristic genomic signatures, and transcriptional profiles. Downregulation of molecules implicated in antigen presentation occurred selectively in highly mutated and rapidly-evolving CRC.

**Conclusions:** These results indicate that CRC carrying alterations in DNA repair pathways display dynamic neoantigen patterns that fluctuate over time. We define CRC subsets characterized by slow and fast evolvability and link this phenotype to downregulation of antigen-presenting cellular mechanisms. Longitudinal monitoring of the neoantigen landscape could be relevant in the context of precision medicine.

## Background

Anticancer therapies based on immune-checkpoint blockade are often remarkably effective but benefit only a minor fraction of cancer patients (1). Several biomarkers of response and resistance to immune modulators have been proposed (2) (3). Among these, the overall mutational burden (number of somatic variants per megabase (Mb)) and the number of predicted neoantigens were highlighted in multiple studies (4) (5) (6). The predictive values of mutational and antigen burdens are still being evaluated in the clinical settings. Both parameters are presently assessed on DNA extracted from individual tissue samples and are typically measured only once in the clinical history of each patient. Alterations in DNA repair pathways, including mutations or promoter hypermethylation of mismatch repair (MMR) effectors (*MLH1*, *MSH2*, etc) or DNA polymerases (polymerase ε and δ) (7) are known to increase the mutational burden and the neoantigen profiles of cancers (8). Whether, and to what extent, neoantigen profiles evolve over time as a result of the inherent genomic instability of individual tumours is largely unknown. We recently reported that in mouse models, inactivation of DNA mismatch repair increases the mutational burden and leads to dynamic mutational profiles resulting in effective cancer immune response (9). Here we exploit CRCs as a model system to understand whether mutational burden and neoantigen profile of human tumours evolve over time as a result of their distinctive genomic landscapes.

## Methods

### CRC cell culture conditions

All cell lines were maintained in their original culturing conditions according to supplier guidelines. Cells were ordinarily supplemented with FBS 10%, 2mM L-glutamine, antibiotics (100U/mL penicillin and 100 mg/mL streptomycin) and grown in a 37°C and 5% CO2 air incubator. To study evolution of cell populations, cell lines were not cloned prior to the experiment or at any subsequent timepoint. Cell lines were thawed in a 10 cm dish. After thaw-recovery, each cell line was screened for the absence of Mycoplasma contamination and checked for its identity, referred below as Quality Control (QC)). To preserve heterogeneity, upon thawing individual lines were expanded to at least 10^8^ cells. At this point for each model, cells were counted, and the percentage of alive/dead cells was calculated. At the beginning of the experiment (T0), 4 × 10^7^ live cells were distributed as follows: (A) 2 × 10^6^ cells were re-plated in a 10cm dish for *in vitro* propagation, (B) 3×10^7^ cells were used for *in vivo* experiments, (C) 2 × 10^6^ cells were frozen, (D) 3 pellets (2×10^6^ cells each) were frozen for DNA, RNA and protein extraction. Cells plated as in A were kept in culture changing medium twice a week and dividing them at constant splitting rate, determined before initiating the experiment. In details, splitting was performed before full confluency was achieved. The number of cells that were split and the number of passages and days of culture were recorded for each cell model to calculate the doubling time. During *in vitro* culture, cell populations were collected at the following pre-determined time points: 30 days (T30), 60 days (T60) and 90 days (T90) days from T0. At each time point a fraction of the cells were put aside (note that this did not affect the rate of passaging described below) and pellets (2×10^6^ each) were collected for DNA, RNA and protein extraction. QC was repeated at each time point.

### Cell quality control (QC)

Cells were screened for absence of Mycoplasma contamination using the Venor®GeM Classic kiy (Minerva biolabs). The identity of each cell line was checked before starting each experiment and after every genomic DNA extraction by PowerPlex® 16 HS System (Promega), through Short Tandem Repeats (STR) at 16 different loci (D5S818, D13S317, D7S820, D16S539, D21S11, vWA, TH01, TPOX, CSF1PO, D18S51, D3S1358, D8S1179, FGA, Penta D, Penta E, and amelogenin). Amplicons from multiplex PCRs were separated by capillary electrophoresis (3730 DNA Analyzer, Applied Biosystems) and analysed using GeneMapper v 3.7 software (Life Technologies).

### Microsatellite instability (MSI) status

The MSI status was assessed with the MSI Analysis System kit (Promega). The analysis requires a multiplex amplification of seven markers including five mononucleotide repeat markers (BAT-25, BAT-26, NR-21, NR-24 and MONO-27) and two pentanucleotide repeat markers (Penta C and Penta D). The products were analysed by capillary electophoresis in a single injection (3730 DNA Analyzer, ABI capillary electrophoresis system (Applied Biosystems). Then the results were analysed using GeneMapper V5.0 software.

### DNA extraction and exome sequencing

Genomic DNA (gDNA) was extracted from CRC cell lines, xenografts, and PDXs using Maxwell® RSC Blood DNA kit (AS1400, Promega). DNA was sent to IntegraGen SA (Evry, France) that performed library preparation, exome capture, sequencing, and data demultiplexing. Final DNA libraries were pair-end sequenced on Illumina HiSeq4000 as paired end 100b reads.

### Mutational analysis in cell lines

When cell lines were passaged in mice or when analysing patient derived xenografts, Fastq files were first processed with Xenome (10) to remove reads of mouse origin. Reads files were aligned to the human reference hg38 using BWA-mem algorithm (11) and then the “rmdup” samtools command was used to remove PCR duplicates (12). On the resulting aligned files, we observed a median depth of 138x with 98% of targeted-region covered by at least one read. Bioinformatic modules previously developed (13) (9) by our laboratory were used to identify single nucleotide variants (SNVs) and indels. The mutational characterization of the 64 cell lines at time point 0 was assessed by calling the alterations against the hg38 reference annotation. Then a series of filters were used to remove germline variants and artifacts: alleles supported by only reads with the same strand, excluding start and end read positions from the count, were discarded; variants called with allelic frequency lower than 10% as well a p-value greater than 0.05 (binomial test calculated on allele count and depth of each sample) were excluded; common dbSNP version 147 and a panel of normal (40 samples) from previous sequencing were used to annotate and filter germline variants and sequencing artifacts. The variant calls of 45 cell lines at time point 90 and the 18 cell lines explanted from mice were performed using the allele comparison strategy between the same cell line at time 0 and time point 90 and xenograft respectively. Only variants present at time point 90 (or in xenograft) were kept. Artifact removal was employed as described above. To calculate the tumour mutational burden (number of variants/Mb), only coding variants were considered. Those variants were used to predict neoantigens using previously published methods (14) (9). Briefly, RNAseq data were used as input of “OptitypePipeline” (15) to assess the HLA status of each samples at time point 0, then NetMHC 4.0 software (16) was employed to analyse mutated peptides derived from variants calls using kmer of 8-11 length. Next, for each SNV we modified the corresponding cDNA in the selected position and we examined the 5’ and 3’ context. The latter was set taking into account the length (in terms of aminoacids) with which the putative antigen could bind HLA. We translated the cDNA and feed mutant peptide to netMHC with the proper HLA(s). For frameshifts, we applied the same approach considering every possible peptide generated by the new frame. Finally, RNAseq data were used to annotate and then filter according to expression values (Fragments Per Kilobase Million (FPKM) > 10). Only predicted neoantigens with a strong binding affinity (Rank < 0.5) were considered for further analysis.

### Mutational analysis of patient-derived xenograft

WES of patient-derived xenografts was performed at IntegraGen SA (Evry, France). Sequenced samples included a microsatellite stable (MSS), a microsatellite unstable (MSI), and a *POLE* mutant case (5, 7, and 6 respectively). Samples were analysed with the same bioinformatic pipeline applied to cell lines and murine reads were first removed using Xenome (10). A median depth of 130x and with 98% of targeted-region covered by at least one read was observed. All 18 PDXs samples were characterized by calling alterations against the hg38 reference annotation. For each generation, with the exception of first one, the mutational evolution was inferred by subtracting the mutations of the previous generation. Second generation samples were compared to the first-generation samples, samples from the 3rd generation were compared to the 2nd generation samples, and so on.

### Ploidy Estimation

Gene copy-number (GCN) was calculated in a two-steps approach: initially we treated the cell line as diploid and considered the median read depth of all coding regions as the level for 2N ploidy. We also calculated the median read depth for every gene. The ratio between the two median values was then considered as the relative GCN. In the second step, to estimate the overall ploidy we segmented all chromosomes using a custom script that implements circular binary segmentation. Finally, we exploited the distribution of allelic frequencies for individual segments to assess the absolute GCN. This was necessary since distinct ploidy levels have different expected distributions. For example, a 2N ploidy status has a bell-shaped curve with a peak on 50%, a 3N ploidy is expected to have two peaks on 33% and 66%, etc.

### Mutational Signature

Mutational signatures were calculated using the web application ‘Mutational Signatures in Cancer’ (MuSiCa) (17). The profile of each signature is calculated using the six substitution subtypes: C>A, C>G, C>T, T>A, T>C, and T>G (all substitutions are referred to by the pyrimidine of the mutated Watson–Crick base pair). Information on nucleotides 5’ and 3’ to each mutated base are incorporated to generate 96 possible mutation types. For each sample, a tab-separated values file was created with chromosome, position, reference, and alternate alleles. Only samples with at least 10 mutations were included. The output file of MuSiCa, that includes the contribution values of 30 signatures (18), was used to create a clustermap with *seaborn*, a *Python* data visualization library, setting Euclidean metric and the average linkage method.

### Doubling time

Cell lines were passaged *in vitro* for a minimum of 85 to a maximum of 103 days. Each passage was performed before full confluency was achieved and the total number of doublings was annotated for each cell model. Two parameters, number of passages (n), and days of culture (t), were used to estimate the growth rate (GR) and the doubling time (DT) assuming that: every division is an independent random event; probability distribution of division is equal for all cells and it is an exponential distribution; the number of cells in each plate before confluence is fixed (K). The growth rate is defined as GR=log_n_(2)÷DT (19). The estimated number of cells at time t is defined as N(t)=N(0)×e^(GR×t)^ where N(0) is the number of cells at time 0. Therefore GR=log_n_(N(t)÷N(0))÷t where N(t)÷N(0)=(K×2^n^)÷(K×2^0^)=2^n^ and so GR=log_n_(2^n^)÷t. Finally, DT=t×log_n_(2)÷ log_n_(2^n^).

### RNA extraction and RNAseq analysis

Total RNA was extracted from a pellet of CRC cells (2×10^6^ cells) using Maxwell® RSC miRNA Tissue Kit (AS1460, Promega), according to the manufacturer’s protocol. The quantification of RNA was performed by Thermo Scientific Nanodrop 1000 (Agilent) and Qubit 3.0 Fluorometer (LifeTechnologies). RNA integrity was evaluated with the Agilent 2100 Bioanalyzer using the Agilent RNA 6000 Nano Kit. Total RNA (800 ng) with RNA integrity number (RIN) score between 9 and 10 was used as input to the Illumina TruSeq RNA Sample Prep Kit v2-Set B (48Rxn), according to the manufacturer’s protocol. The standard RNA fragmentation profile was used (94 °C for 8 min for the TruSeq RNA Sample Prep Kit). PCR-amplified RNA-seq library quality was assessed using the Agilent DNA 1000 kit on the Agilent 2100 BioAnalyzer and quantified using Qubit 3.0 Fluorometer (LifeTechnologies). Libraries were diluted to 10 nM using Tris-HCl (10 mM pH 8.5) and then pooled together. Diluted pools were denatured according to standard Illumina protocol and 1.8 pM were run on NextSeq500 using high output Reagent cartridge V2 for 150 cycles. A single-read 150-cycle run was performed. FastQ files produced by Illumina NextSeq500 were aligned using MapSplice2 (20) transcriptome-aware aligner using hg38 assembly as reference genome. The resulting BAM files were post-processed to translate genomic coordinates to transcriptomic ones and to filter out alignments carrying insertions or deletions (which RSEM does not support) or falling outside the transcriptome regions. The post-processed BAM alignment was given as input to RSEM (21) for gene expression quantification using GENCODE v22 as gene annotation.

### Differential expression analysis

The abundance quantification generated with RSEM provides the FPKM and the expected counts for each gene. The latter was used to perform genes differential expression analysis with DESeq2 R package (library Bioconductor) (22) given two distinct groups of interest, one of which considered as the reference. Genes were considered as differentially expressed if the adjusted p-value was less than 0.05, and the log2 fold change was less or equal to −1 (if median FPKM value of the reference group was greater or equal to 10) or the log2 fold change was greater or equal to 1 (if median FPKM of the target group was greater or equal to 10). The analyses were performed between the following groups: MSI *vs* MSS (reference); hypermutated *vs* non-hypermutated (reference); “EVOLVING-CRC” *vs* “STABLE-CRC” (reference). Hypermutated group included MSI and MSS POLE mutated cell lines (18 samples). “EVOLVING-CRC” group included all samples with at least 10 alterations acquired per day. A multi-factor configuration of the expression analysis was designed including extra variables of interest such as growth rates or the number of mutations normalized to doubling time.

### Pathway analysis

Genes differentially expressed were then analysed with g:Profiler (23), an online pathway analysis tool that takes a list of genes and assigns them to different families of biological functions. We set the query options to select significant biological processes only and we retained (for further analysis) only the top most families of the hierarchy (depth 1).

### Xenograft mouse model

Each CRC cell line (5×10^6^ cells) was injected subcutaneously into both flanks of two 6-week-old female NOD (nonobese diabetic)/SCID (severe combined immunodeficient) mice (Charles River Laboratory). Tumour size was measured twice a week and calculated using the formula: *V* = ((d)^2^ × (D)) ÷2 (d = minor tumour axis; D = major tumour axis). Tumours were explanted when they reached a volume of 1000mm^3^. The investigators were not blinded, and measurements were acquired before the identification of the cages.

### Patient-derived mouse model

Tissue from hepatic metastasectomy of CRC patients was collected at surgery and implanted in NOD-SCID mice as described previously (24). When reaching a volume of 1500-2000mm^3^, the tumours were explanted, fragmented, and serially passaged in new mice. At each passage, part of the material was frozen for molecular analyses. Samples’ genetic identity was determined by Sequenom-based analysis of 24 highly variable SNPs of germline DNA (Table 5), confirmed by analysing pre-implantation tumour material, and then validated every second passage in mice. The study population consisted of matched tumour and normal samples from 3 CRC patients that underwent surgical resection of liver metastases at the Candiolo Cancer Institute (Candiolo, Torino, Italy) and at the Mauriziano Umberto I Hospital (Torino) between 2009 and 2013. Patients signed informed consent, and the study was approved by the relevant institutional Ethics Committees.

### Western blotting analysis

Proteins were extracted by solubilizing the cells in boiling SDS buffer (50 mM Tris-HCl [pH 7.5], 150 mM NaCl, and 1% SDS). Samples were boiled for 5 minutes at 95°C and sonicated for 10 seconds. Extracts were clarified by centrifugation, normalized with the BCA Protein Assay Reagent kit (Thermo). Equal amounts of proteins (20μg) were loaded in each lane. Proteins were separated by PAGE and transferred to nitrocellulose sheets. Western blot detection was performed with enhanced chemiluminescence system (GE Healthcare) and peroxidase conjugated secondary antibodies (Amersham). The following primary antibodies were used for western blotting: anti-beta2 Microglobulin [EP2978Y] (ab75853, Abcam), anti-MLH1 (ab92312, Abcam), anti-MSH2 (ab70270), Abcam), anti-MSH6 [EPR3945],(ab92471, Abcam), anti-MSH3 PA527864, Invitrogen, anti-PMS2 EPR3947 (Cell Marque Corporation, USA), anti-actin (I-19) (sc1616, Santa Cruz), anti-HSP 90α/β (H-114, sc-7947, Santa Cruz). Images were acquired with Chemidoc (Biorad) and western blot band intensity was analysed using Image Lab software (Biorad).

## Results

We selected from our database 64 CRC cell lines designed to recapitulate clinically relevant characteristics of CRC patients (Table 1 and Suppl. Fig. 1a). Whole exome sequencing and RNAseq were performed on all models. Using previously developed computational tools and bioinformatic algorithms (13) (25) (14) (26), we measured mutational burden (alterations per Mb) assessing both SNVs and frameshifts (Fig. 1a,b) Scrutiny of genomic alterations highlighted that MSI cell lines and those carrying known *POLE* hotspot mutations had higher number of mutations per Mb as compared to MSS cell lines (Fig. 1a). The type of DNA repair alterations occurring in each model affected the nature of mutations: MSI cells displayed higher number of frameshifts and indels than *POLE* mutant cell lines, the opposite was true for SNVs (Fig. 1c, d).

**Fig. 1.**
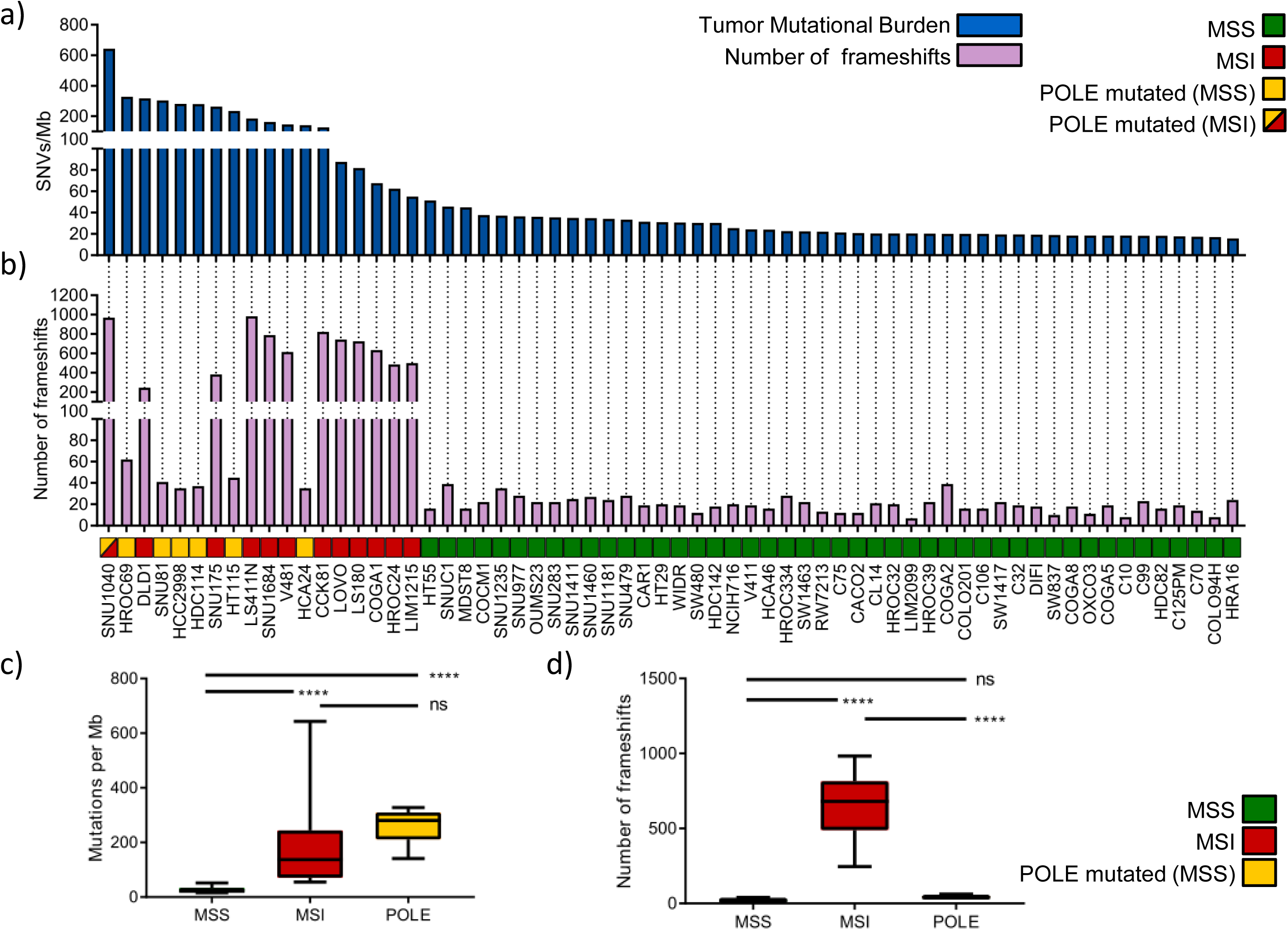
Analysis of mutational burden in a panel of 64 CRC cell lines. Mutational characterization and comparison of SNVs and frameshifts among MSS (46 samples), MSI (12 samples) and POLE mutated (6 samples) of CRC models. a) The distribution of SNVs per Mb of coding DNA at time 0 is shown for each cell line. b) The number of frameshift mutations at time 0 is shown for each cell line. c) The number of SNVs per each group is shown (‘MSS’ refers to MSS cells without POLE mutations; ‘MSI’ includes MSI cells, as well as the SNU1040 cell line which is both MSI and POLE mutated; ‘POLE’ includes only MSS cell lines carrying a POLE mutation). d) The number of frameshifts per group is shown. The center line of each box plot indicates the median. P < 0.0001.

**Table 1.** Molecular and functional characteristics of the indicated cell lines.

| **SAMPLE** | **Microsatellites Status** | **Altered MSI Markers** | **Genome Evolvability** | **Split ratio** | ***In vitro* Doubling Time** | **Growth rate** |
| --- | --- | --- | --- | --- | --- | --- |
| C10 | Stable | - | Stable | 0.34 | 2.33 | 0.30 |
| C106 | Stable | - | Stable | 0.35 | 2.50 | 0.28 |
| C125PM | Stable | - | Stable | 0.34 | 2.37 | 0.29 |
| C32 | Stable | - | Stable | 0.18 | 1.54 | 0.45 |
| C70 | Stable | - | Stable | 0.4 | 2.76 | 0.25 |
| C75 | Stable | - | Stable | 0.36 | 2.46 | 0.28 |
| C99 | Stable | - | NA | NA | NA | NA |
| CACO2 | Stable | - | Stable | 0.26 | 1.83 | 0.38 |
| CAR1 | Stable | - | Stable | 0.36 | 2.47 | 0.28 |
| CCK81 | Unstable | bat26-nr21-bat25-mono27-nr24 | Evolving | 0.34 | 2.53 | 0.27 |
| CL14 | Stable | - | NA | NA | NA | NA |
| COCM1 | Stable | - | Evolving | 0.3 | 2.34 | 0.30 |
| COGA1 | Unstable | bat26-mono27-nr24 | NA | NA | NA | NA |
| COGA2 | Stable | - | Stable | 0.32 | 2.22 | 0.31 |
| COGA5 | Stable | - | Stable | 0.20 | 1.69 | 0.41 |
| COGA8 | Stable | - | Stable | 0.22 | 1.67 | 0.41 |
| COLO201 | Stable | - | NA | NA | NA | NA |
| COLO94H | Stable | - | Stable | 0.45 | 2.81 | 0.25 |
| DIFI | Stable | - | NA | NA | NA | NA |
| DLD1 | Unstable | bat26-nr21-bat25-mono27-nr24 | Evolving | 0.07 | 0.98 | 0.71 |
| HCA24 | Stable | - | Evolving | 0.33 | 2.22 | 0.31 |
| HCA46 | Stable | - | Stable | 0.4 | 2.41 | 0.29 |
| HCC2998 | Stable | - | Stable | 0.34 | 2.33 | 0.30 |
| HDC114 | Stable | - | Evolving | 0.23 | 1.69 | 0.41 |
| HDC142 | Stable | - | Evolving | 0.35 | 2.42 | 0.29 |
| HDC82 | Stable | - | NA | NA | NA | NA |
| HRA16 | Stable | - | NA | NA | NA | NA |
| HROC24 | Unstable | bat26-nr21-bat25-mono27-nr24 | Evolving | 0.16 | 1.23 | 0.56 |
| HROC32 | Stable | - | Stable | 0.62 | 6.07 | 0.11 |
| HROC334 | Stable | - | Stable | 0.45 | 2.83 | 0.24 |
| HROC39 | Stable | - | Stable | 0.5 | 3.92 | 0.18 |
| HROC69 | Stable | - | Stable | 0.33 | 2.26 | 0.31 |
| HT115 | Stable | - | Evolving | 0.24 | 1.78 | 0.39 |
| HT29 | Stable | - | Stable | 0.16 | 1.38 | 0.50 |
| HT55 | Stable | - | Stable | 0.33 | 2.21 | 0.31 |
| LIM1215 | Unstable | bat26-nr21-bat25-mono27-nr24 | Evolving | 0.15 | 1.24 | 0.56 |
| LIM2099 | Stable | - | Stable | 0.33 | 2.26 | 0.31 |
| LOVO | Unstable | nr21-bat25-mono27-nr24 | NA | NA | NA | NA |
| LS180 | Unstable | bat26-nr21-bat25-mono27-nr24 | Evolving | 0.25 | 1.90 | 0.37 |
| LS411N | Unstable | bat26-nr21-bat25-mono27-nr24 | Evolving | 0.32 | 2.29 | 0.30 |
| MDST8 | Stable | - | Stable | 0.15 | 1.31 | 0.53 |
| NCIH716 | Stable | - | NA | NA | NA | NA |
| OUMS23 | Stable | - | Stable | 0.26 | 1.70 | 0.41 |
| OXCO3 | Stable | - | Stable | 0.23 | 1.76 | 0.39 |
| RW7213 | Stable | - | Stable | 0.4 | 2.60 | 0.27 |
| SNU1040 | Unstable | bat26-nr21-bat25-mono27-nr24 | Evolving | 0.54 | 4.14 | 0.17 |
| SNU1181 | Stable | - | Stable | 0.62 | 4.65 | 0.15 |
| SNU1235 | Stable | - | Evolving | 0.35 | 2.33 | 0.30 |
| SNU1411 | Stable | - | Evolving | 0.44 | 2.76 | 0.25 |
| SNU1460 | Stable | - | NA | NA | NA | NA |
| SNU1684 | Unstable | bat26-nr21-bat25-mono27-nr24 | NA | NA | NA | NA |
| SNU175 | Unstable | bat26-nr21-bat25-mono27 | NA | NA | NA | NA |
| SNU283 | Stable | - | NA | NA | NA | NA |
| SNU479 | Stable | - | NA | NA | NA | NA |
| SNU81 | Stable | - | Evolving | 0.42 | 2.33 | 0.30 |
| SNU977 | Stable | - | Stable | 0.42 | 2.71 | 0.26 |
| SNUC1 | Stable | - | NA | NA | NA | NA |
| SW1417 | Stable | - | NA | NA | NA | NA |
| SW1463 | Stable | - | NA | NA | NA | NA |
| SW480 | Stable | - | Stable | 0.18 | 1.64 | 0.42 |
| SW837 | Stable | - | Stable | 0.27 | 2.07 | 0.34 |
| V411 | Stable | - | Stable | 0.22 | 1.91 | 0.36 |
| V481 | Unstable | bat26-nr21-bat25-mono27-nr24 | NA | NA | NA | NA |
| WIDR | Stable | - | NA | NA | NA | NA |

Alterations in MMR and *POLE* genes are listed in Table 2 and Suppl. Fig. 1b. The cell line with the highest number of variants (SNU1040) carried inactivating alterations in both *MLH1* and *POLE* (Suppl. Fig. 1b). Altogether, these results are consistent with what has been reported in CRC patients carrying alterations in the MMR DNA repair pathway, indicating that the cell models included in this study broadly recapitulate what is observed in clinical specimens (27).

**Table 2.**
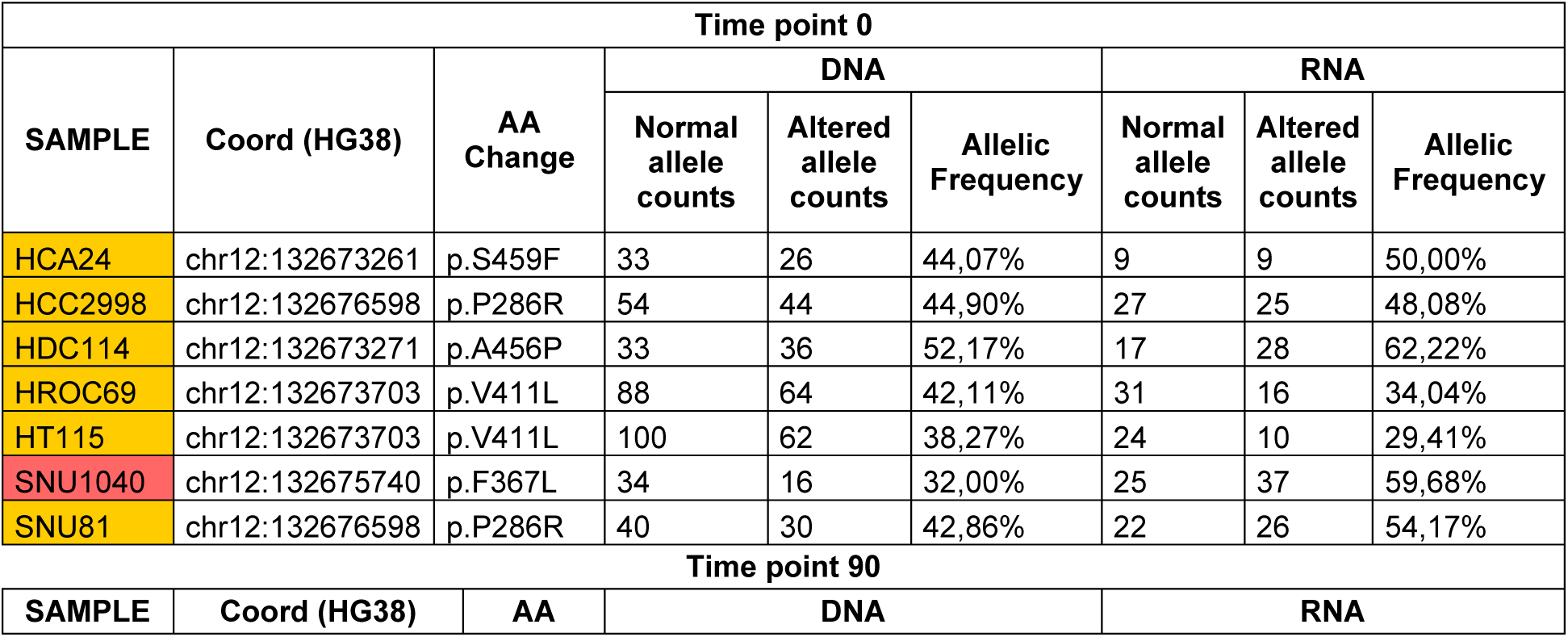
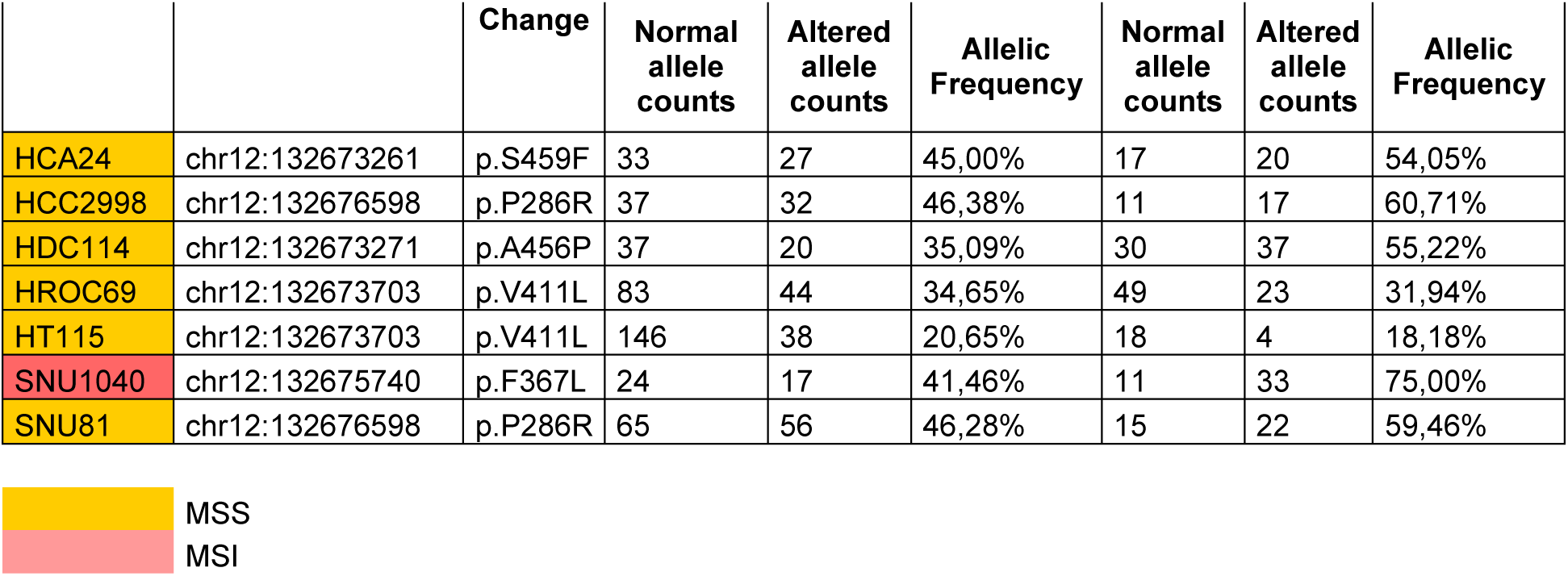
POLE mutations in CRC cells.

To assess whether, and to what extent the basal mutational profiles (Time 0: T0) evolved over time, we passaged 45 cell lines for 90 days and collected a second set of samples (Time 90: T90) (Suppl. Fig. 2). These were subjected to WES and analysed using the computational pipeline described above. Across all cell lines globally, the total mutational burden was similar between T0 and T90 (Suppl. Fig. 3). However, when the T0 and T90 mutational profiles were compared, prominent differences were detected among models sharing specific DNA repair defects (Fig. 2a). Specifically, the mutational landscapes of most MSI and *POLE* mutant cells evolved very rapidly through the generation of novel SNVs and frameshifts (Fig. 2a). On the contrary, the majority of MSS models showed more stable profiles (Fig. 2a). We sought to minimize confounding effects due to differences in cell-intrinsic doubling times (Table 1), we therefore calculated the doubling time of all cell models (Table 1, Suppl. Fig. 4). Notably, evolvability trends remained apparent after normalization for doubling time (Suppl. Fig. 5). We designated rapidly evolving CRC cells as “EVOLVING-CRC” and evolutionary stable CRC cell as “STABLE-CRC” (Table 1).

**Fig. 2.**
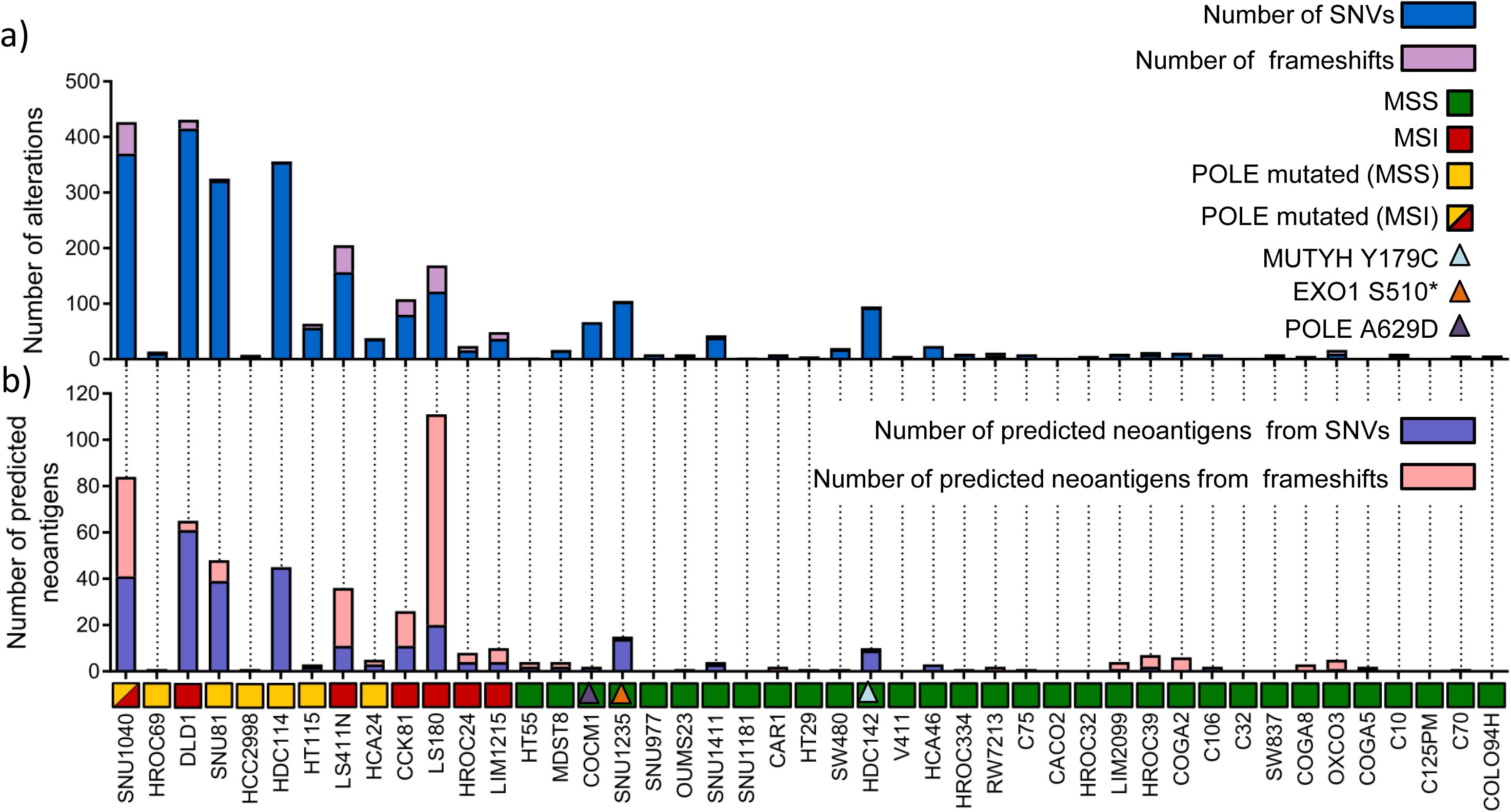
*In vitro* evolution of mutational landscape in 45 CRC cell lines. Mutational characterization of CRC cells after 90 days of culture (T90*) in vitro*. a) Bar charts show the number of novel alterations (SNVs and frameshifts) acquired at T90 (not present at T0) for each cell line. b) The number of predicted neoantigens (see Methods) is shown. Each bar represents putative neoepitopes derived from SNVs and frameshifts.

We empirically define EVOLVING-CRCs as those cells that acquire 10 alterations (or more) per day after normalizing mutation data to the doubling time of cell lines (Table 1). Moreover, EVOLVING-CRCs often carried alterations in multiple genes involved in distinct DNA repair functions, suggesting that defects in several DNA damage response pathways might be co-selected (Suppl. Fig. 1b). The expression of MMR genes was assessed by western blot at T0 and T90 and no differences were observed (Suppl. Fig. 6).

The genome of four CRC lines classified as MSS (SNU1235, COCM1, HDC142, and SNU1411) exhibited dynamic mutational profiles (Fig. 2). In an attempt to decipher the molecular basis of these findings, whole exome data of the outliers were carefully examined, focusing on genes previously implicated in DNA repair pathways that are not routinely subjected to scrutiny in CRC patients. We found that SNU1235 and HDC142 models carried biallelic alterations in the *EXO1* (S510*) and *MUTYH* (S179C) genes, respectively. The exonuclease EXO1 is implicated in both MMR (it binds MLH1) and base excision repair (28), while *MUTYH* encodes a DNA glycosylase that is involved in oxidative DNA damage repair and is part of the base excision repair pathway (29). Germline mutations in *MUTYH* cause MUTYH Associated Polyposis (MAP) (30). Scrutiny of the COCM1 exome revealed a *POLE* variant (A629D). A629 is localized in a region of *POLE* highly conserved during evolution (Suppl. Fig. 7). The A629D change is potentially damaging according to the SIFT (31) and Polyphen (32) algorithms, which predict the putative impact of amino acid substitutions on human proteins using structural and comparative evolutionary considerations.

We next addressed how longitudinal evolution of CRC cell genomes affected their predicted neoantigen profile. To this end, WES, RNAseq, and HLA prediction data were combined as previously described (9). In detail, we identified genomic variants that satisfied three criteria: (i) emerged over time, (ii) occurred in transcribed genes, and (iii) scored positively when HLA I matching algorithms were applied. The variants that emerged after deploying the above computational pipeline were classified as putative neoantigens (Fig. 2b). Hypermutated and “EVOLVING-CRC” cells displayed higher levels of putative neoantigens compared to slowly evolving CRC cells (Fig. 2b). Moreover, and consistent with their predicted effects on antigenicity, a high prevalence of indels and associated frameshifts, which occur in MSI CRCs, translated into higher numbers of predicted neoantigens in this subset (Fig. 2b).

Next, we studied whether in parallel to mutation’ gains we could also detect loss of variants over time. For this reason, we tracked lost and gained alteration in ‘evolving’ cell lines over time. As expected, variants that did not change over time showed high allelic frequency, likely reflecting their clonal (trunk) status. Mutations that emerged or were lost showed lower allelic frequency (Fig. 3).

**Fig. 3.**
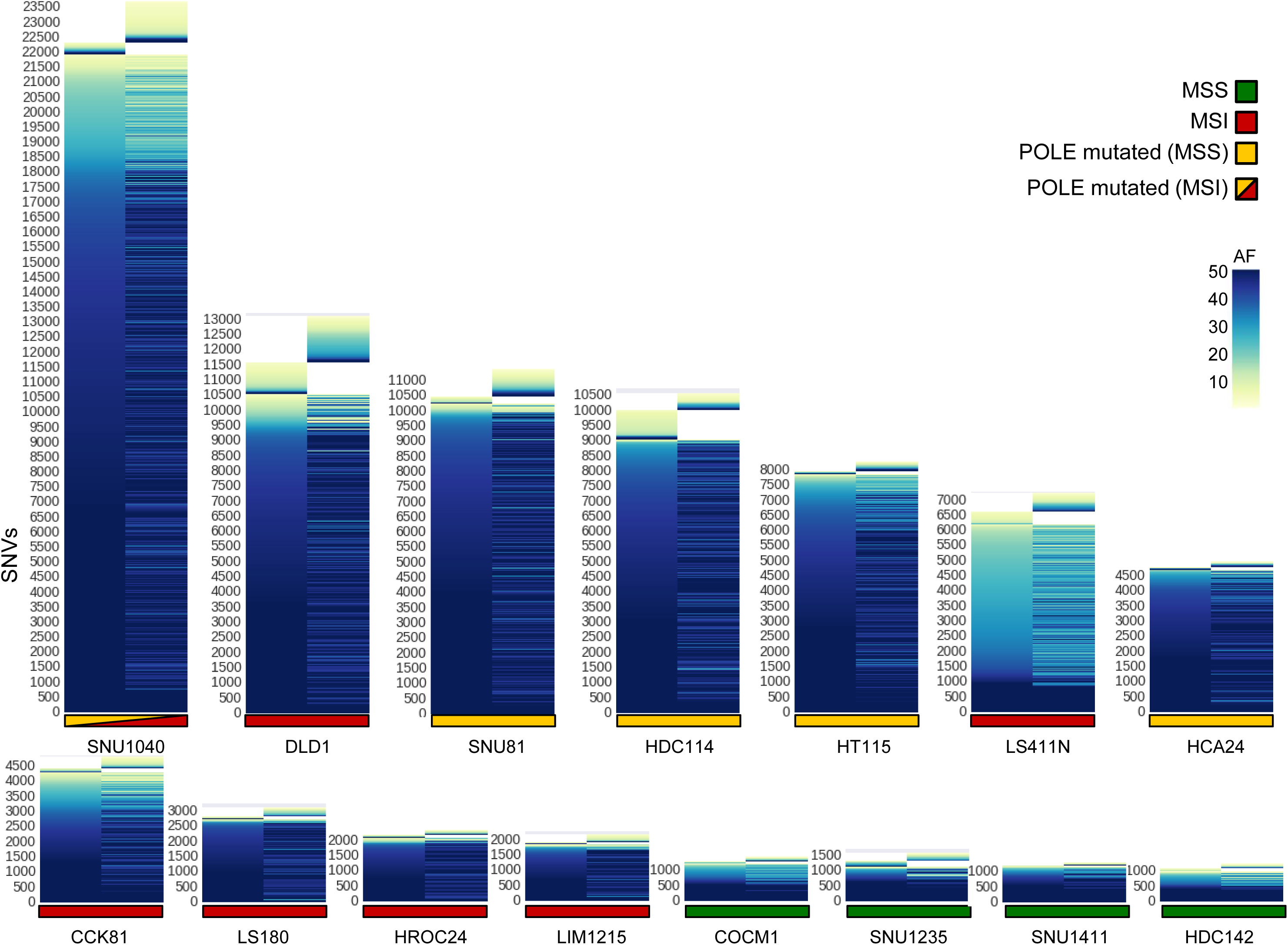
Lost and gained mutations across ‘evolving’ CRC cell lines. For each CRC model the allelic frequency of SNVs at T0 and T90 are shown. Mutations were called against the reference genome (hg38) with allelic frequency > 1. The y-axis reports all the mutations found in each cell line, whereas the time points data are reported on x-axis.

Mutational signatures are characteristic combinations of mutation types arising from mutagenesis processes such as alterations in DNA replication, exposure to DNA damaging agents, tissue culture conditions, and DNA enzymatic editing (18). In human tumours, over 30 mutational signatures have been identified, a subset of which are linked to defective DNA repair pathways. For example, signatures 6, 15, 20, and 26 are associated with MMR defects, signature 10 is linked to inactivating mutation in the proofreading domain of DNA polymerases, while signature 18 appears to explain the rise of 8-oxoG:A mismatches due to *MUTYH* biallelic alteration (33).

We reasoned that the remarkable evolvability observed in a subset of CRC cells might be reflected in their mutational signatures. To test this, we first identified mutational signatures at T0. As expected, MSI cells displayed signatures 6, 15, 20, and 26, while *POLE* mutant cells showed primarily mutational signature 10 (Suppl. Fig. 8).

We next assessed which signatures were acquired (remained active) during replication of the cells *in vitro* by comparing samples collected at T0 and T90. We found that in most instances, DNA alterations linked to MMR and *POLE* defects continued to occur over time, indicating that the corresponding DNA repair capabilities were permanently disabled (Fig. 4a).

**Fig. 4.**
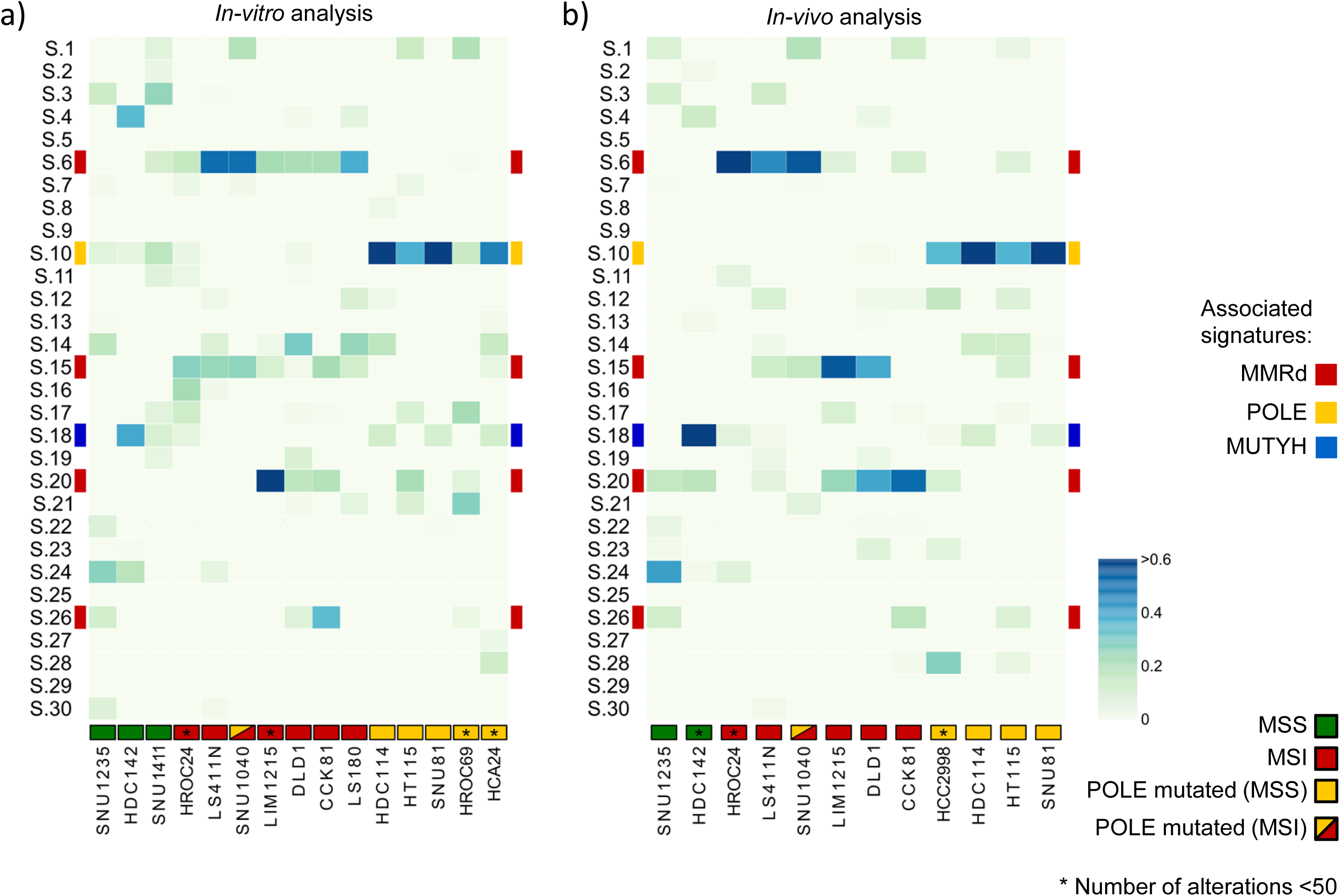
Mutational signatures associated with alterations emerging during *in vitro* or *in vivo* CRC propagation. Analysis of 30 validated cancer associated mutational signatures in hypermutated/rapidly evolving CRC cell lines. Signatures associated to MMR-deficient (6, 15, 20 and 26), *POLE*-dependent (10) and *MUTYH*-associated polyposis (18) are highlighted. Analysis and clustering were performed as reported in Methods. a) Heatmap of signature contributions during replication of CRC cells *in vitro* by analysing alterations acquired at T90. b) Heatmap of signature contributions during replication of the CRC cells *in vivo* by comparing xenograft tumours to the corresponding cells at T0 (See Methods for detailed information).

Replication of cancer cell populations in 2D is thought to encounter little or no selective pressure as the cells are cultured in the same conditions for many generations before the experiment is started. To monitor mutational and neoantigen evolution under more stressful (selective) conditions, CRC cells including MSS, MSI, and *POLE* models were transplanted in immunodeficient (NOD SCID) mice and allowed to grow until they reached approximately 1000mm^3^ in size, after which tumours were excised. Although NOD SCID mice have no adaptive immunity, the mouse stromal microenvironment and elements of cellular innate immunity are known to affect the growth of human cancer cells *in vivo* (34). DNA samples were obtained before implantation and at the end of the experiment. WES was performed, and the data were analysed with the same bioinformatic pipeline applied to cells grown *in vitro*. The mutational profiles revealed higher evolutionary rates *in vivo* than *in vitro* (Suppl. Fig. 9a,b). This translated in increased levels of predicted neoantigens *in vivo*, (Suppl Fig 9c). Notably, mutational signatures linked to MSI status and *POLE* mutations were more marked *in vivo* than *in vitro* (Fig. 4b, Suppl Fig 10).

Next, we asked whether the evolutionary trajectories observed in CRC cells with alterations in DNA repair pathways also occurred in human CRC with analogous molecular profiles. To this end, we selected MMR proficient, MMR deficient, and POLE mutant cases (Table 3) from our extensive patient-derived CRC xenograft biobank (35). Each model was serially transplanted for at least four generations in immunodeficient mice as described in the phylogenetic tree (Fig. 5a). Samples collected at each transplantation were subjected to WES. In some instances, simultaneous transplantation of the same tumour in two animals allowed acquisition of independent measurements for each generation. NGS data were analysed with the bioinformatic pipeline applied to cells grown *in vitro*. These experiments revealed remarkable differences in the evolvability of MSS, MSI, and *POLE* CRC models *in vivo*, and indicated that these characteristics also occurred in patient-derived CRC samples (Fig. 5b, c). As expected, high frequency (clonal-trunk) variants were conserved across generations. Interestingly, the *in vivo* results differ from those obtained in cell models *in vitro*. We find that in PDX models, not only sub-clonal but also clonal populations can emerge in subsequent generation of colorectal cancers with DNA repair defects (Fig 6).

**Table 3.**
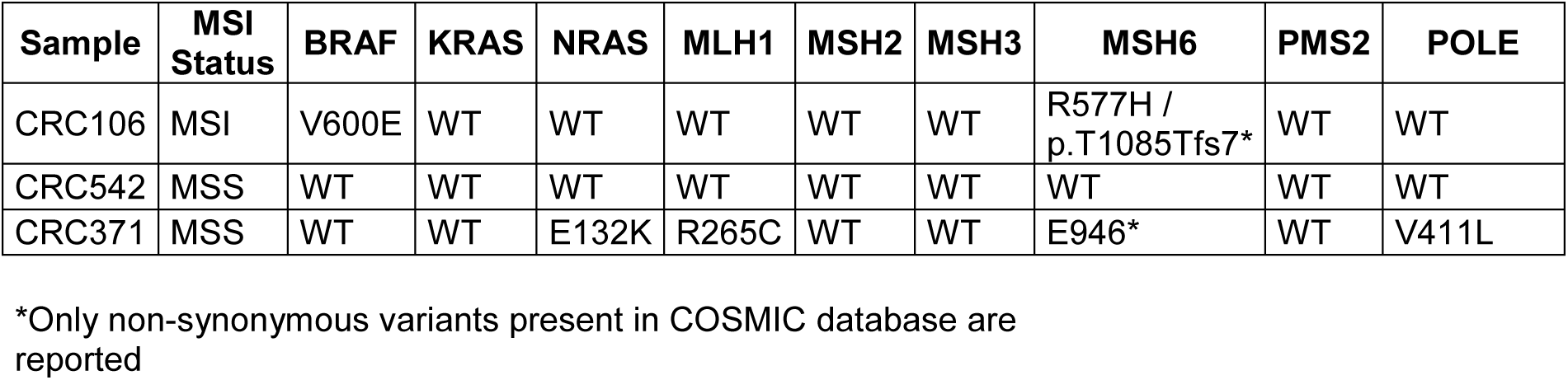
Molecular characterization of patient-derived xenografts.

**Fig. 5.**
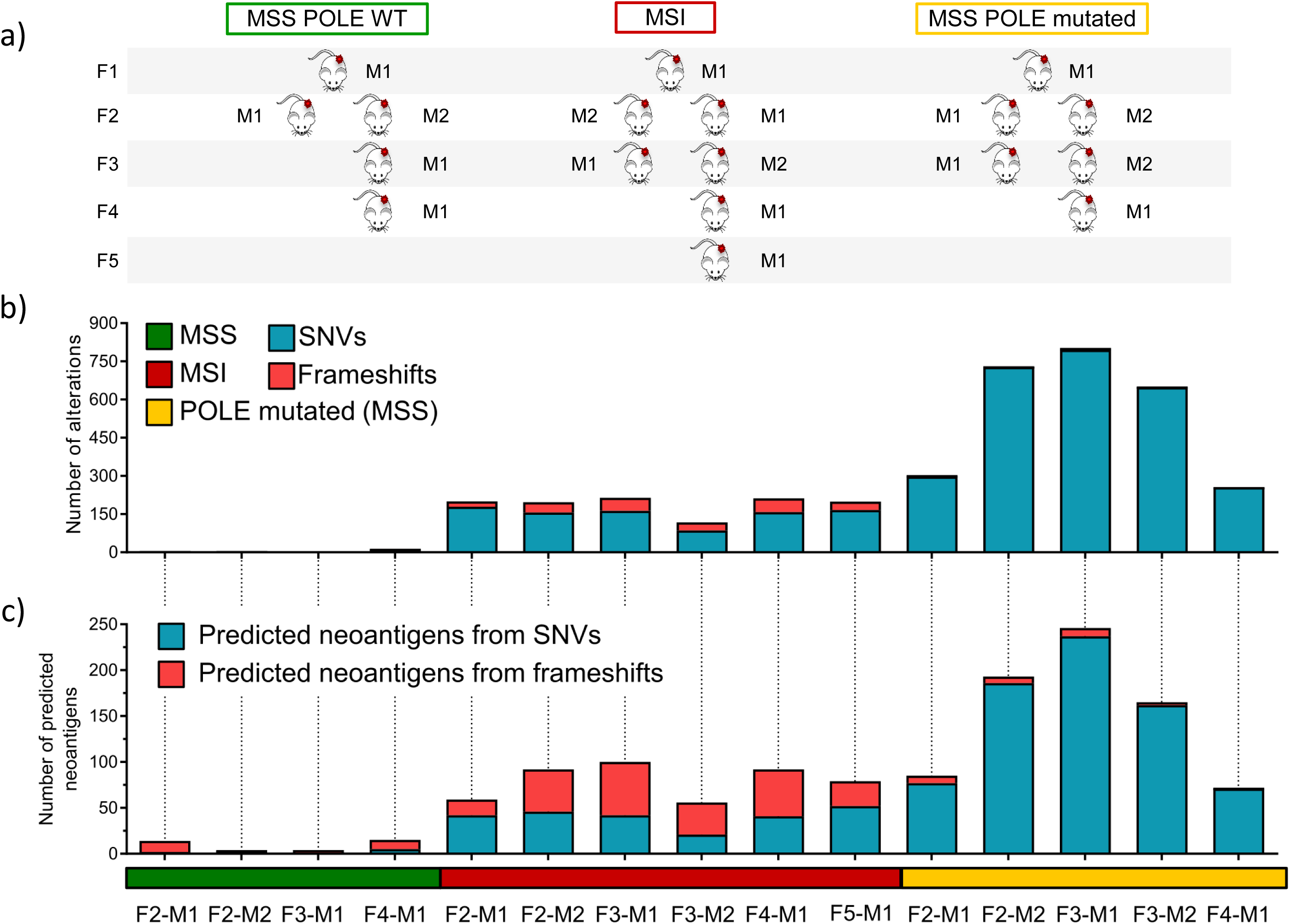
Genomic evolution in patient-derived xenografts. Phylogeny of the indicated patient-derived xenograft and their molecular characterization. a) MSS, MSI and *POLE* mutant samples were serially transplanted for at least four generations (F1-F4) in NOD/SCID mice as shown. Samples collected at each passage were subjected to WES. b) WES data of each generation were compared with those obtained from the previous generation. Bar graphs show *de novo* acquired SNVs and frameshifts at each generation. c) The number of predicted neoantigens in each PDX is shown. Each bar represents putative neoepitopes derived from SNVs and frameshifts. (See Methods for detailed information).

**Fig. 6.**
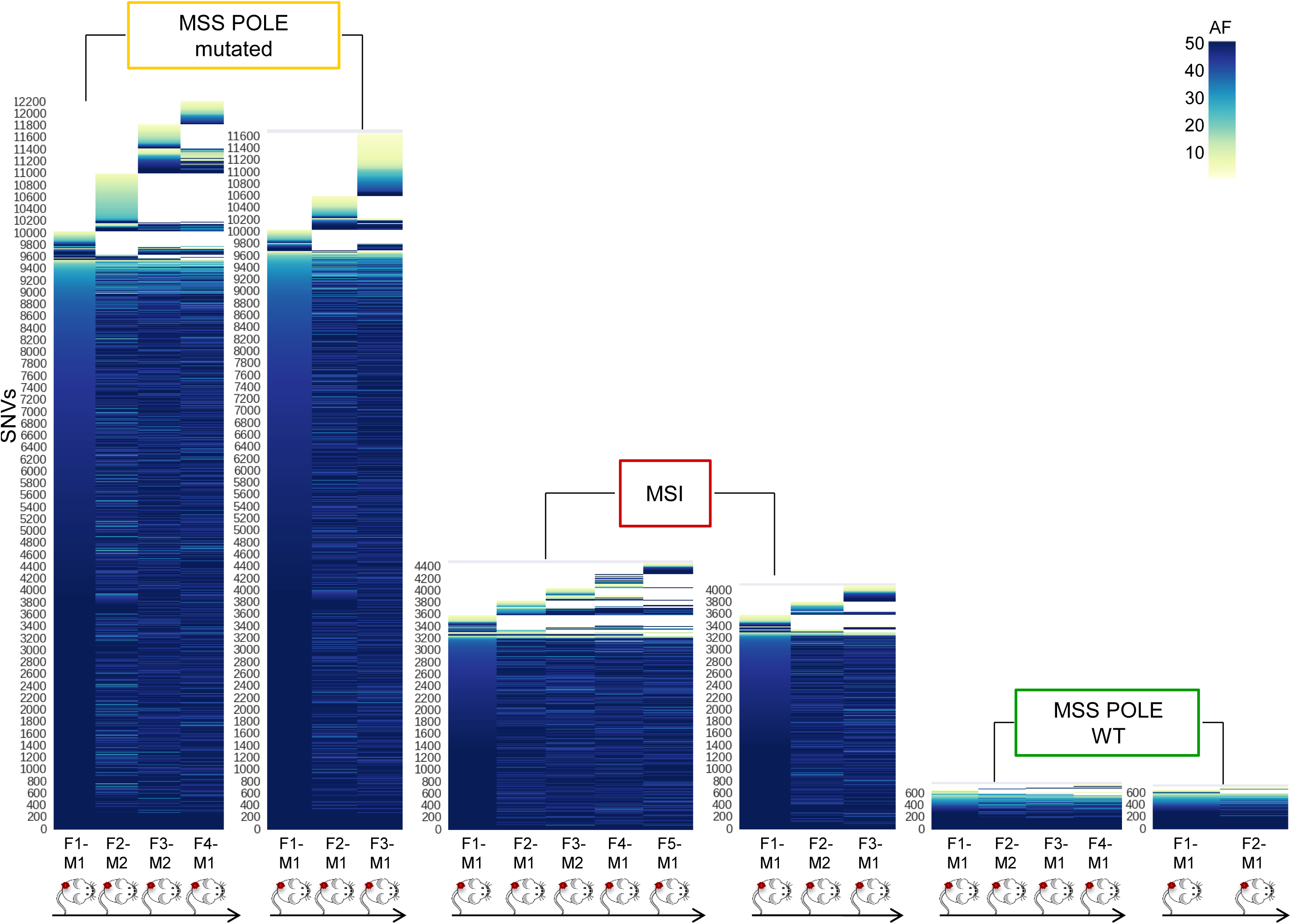
Lost and gained mutations across the indicated PDX generations. The color code defines allelic frequencies of acquired SNVs at each generation (with allelic frequency > 1). The y-axis lists all SNV identified in each branch, the mouse generation (genealogy) are reported on the x-axis.

Furthermore, in MSI and *POLE* patient-derived xenografts, the mutational signatures were continuously (re)generated and could be clearly recognized (Suppl. Fig. 11 and 12). In non-mutator (slow evolving) cell lines very few mutations emerged over time, and thus the possibility to assess mutational signatures was limited. Because of this, in the slowly-evolving models we were unable to reliably generate mutational signatures.

Distinct subsets of CRCs can be recognized based on histological characteristics, as well as their genomic, epigenetic, and transcriptional profiles. As a result, CRC can be classified into specific subsets, which are often correlated with divergent clinical outcomes (36) (37). The rate of genomic evolution and the dynamics of neoantigen profile have not yet been systematically explored as a method to classify CRC. We therefore asked whether any molecular traits (beyond alterations in DNA repair genes) could distinguish “EVOLVING-CRC” and “STABLE-CRC”. To address this question, we performed unbiased gene copy and transcriptional comparative analyses of CRC cell lines. As previously reported, MSI CRC cells typically carried a close to diploid chromosomal status, while MSS showed elevated aneuploidy (Fig. 7) (38). Interestingly, the most rapidly evolving *POLE* mutant lines, SNU81 and HDC114, also displayed a diploid prevalent phenotype. Nonetheless, copy number and ploidy status could not distinguish “EVOLVING” and “STABLE” CRC models.

**Fig. 7.**
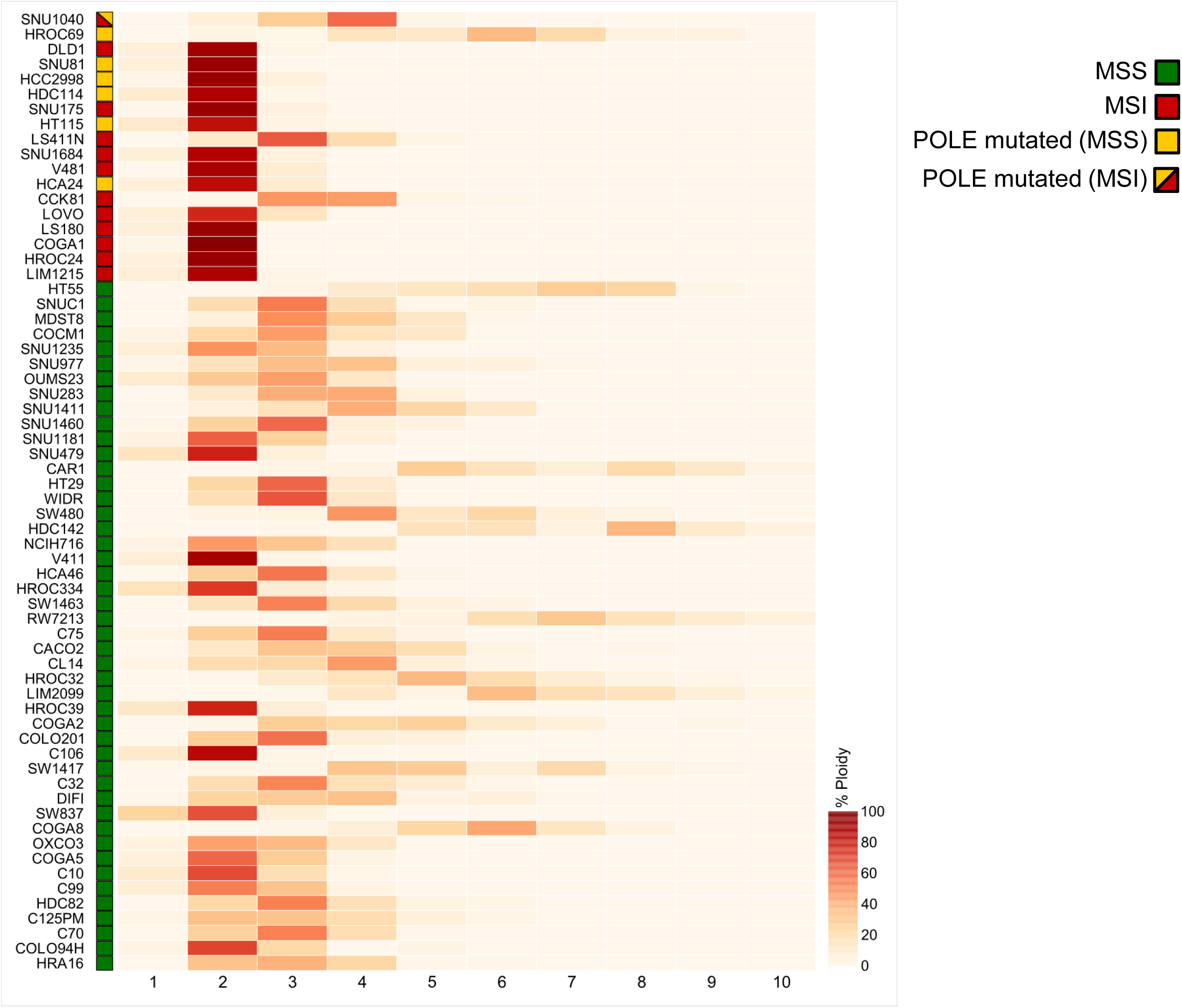
Analysis of cell ploidy in a panel of 64 CRC cell lines. Heatmap showing distribution of ploidy for every segmented region in each cell line. Samples are sorted from most to less mutated as reported in Fig. 1. The percentage (ploidy) is calculated as described in detail in the Methods.

Next, we performed RNAseq on the entire dataset to explore whether transcriptional profiles could classify rapidly evolving CRC lines. Differential analysis of RNAseq data was initially performed comparing the MSS and MSI sample groups. The list of differentially expressed genes was consistent to results previously reported in this setting, and 168 genes were differentially expressed between these two groups (Table 4) (39). Next, we evaluated genes differentially expressed in hypermutated versus non-hypermutated cells, grouping together MSI and *POLE* mutated cell lines and comparing them to the MSS lines (Fig. 8a). Notably, proteins associated with immune response and predominantly with antigen-presenting and antigen recognition functions were consistently downregulated in cell lines with high mutational burden (Fig. 8b). Next, we compared “EVOLVING” and “STABLE” CRC models. The number of genes differentially expressed with significant p-value was smaller due to the reduced number of available samples (Fig. 9a). Beta-2 microglobulin (B2M) was downregulated in most “EVOLVING” as compared to “STABLE” CRCs (Fig. 9b, c). Downregulation of B2M was confirmed at the protein level (Fig. 9c) and was frequently associated with premature stop codons in the *B2M* gene (Fig. 9d). Interestingly, the four MSS models (COCM1, SNU1235, SNU1411, and HDC142) with low mutational burden but dynamic mutational profile also displayed low levels of B2M (Fig.9b, c). Comparison of “EVOLVING” and “STABLE” CRC models pinpointed other genes differentially expressed including CPNE1, IRF1, and PMSB10. These genes are also involved in immune-related processes and their downregulation might similarly reduce immune surveillance of “EVOLVING” CRCs (Fig. 9a and Suppl. Fig. 13). We next performed the analysis showed in Fig 9a in a multivariate fashion taking into account the growth rates of the cells or the number of mutations normalized to the doubling time. The number of statistically significant genes in the multivariate analyses (Suppl Fig 14) was lower but consistent with the findings of Fig 9a. In the future it would be interesting to assess whether the differential expression of genes in fast evolving CRC models has a functional impact. This aspect cannot be causally predicted at this stage.

**Table 4.**
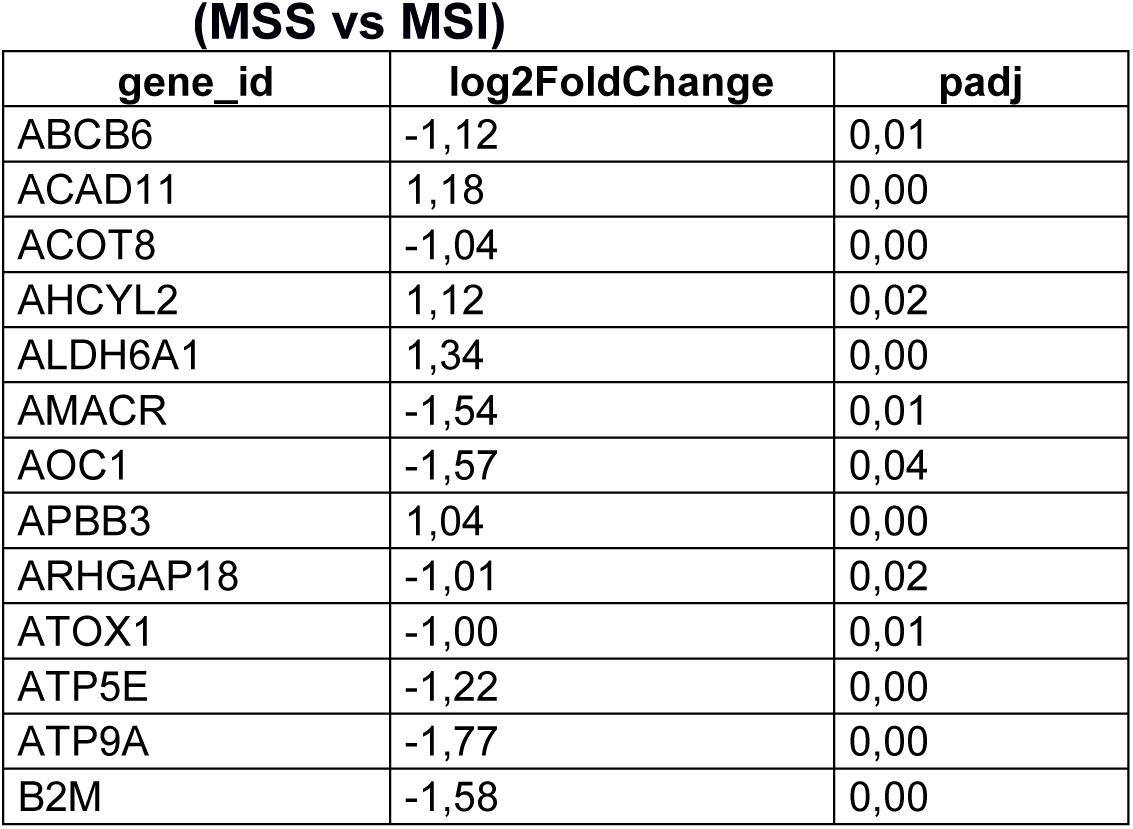
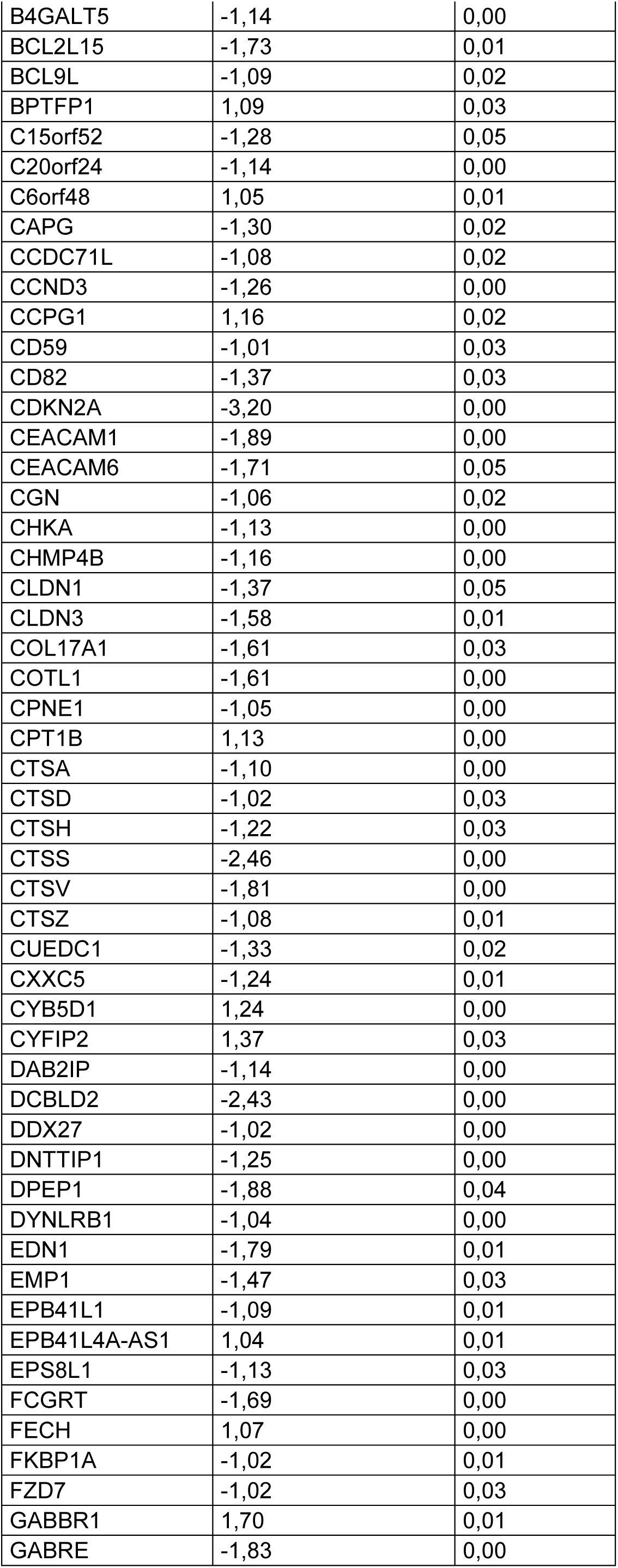
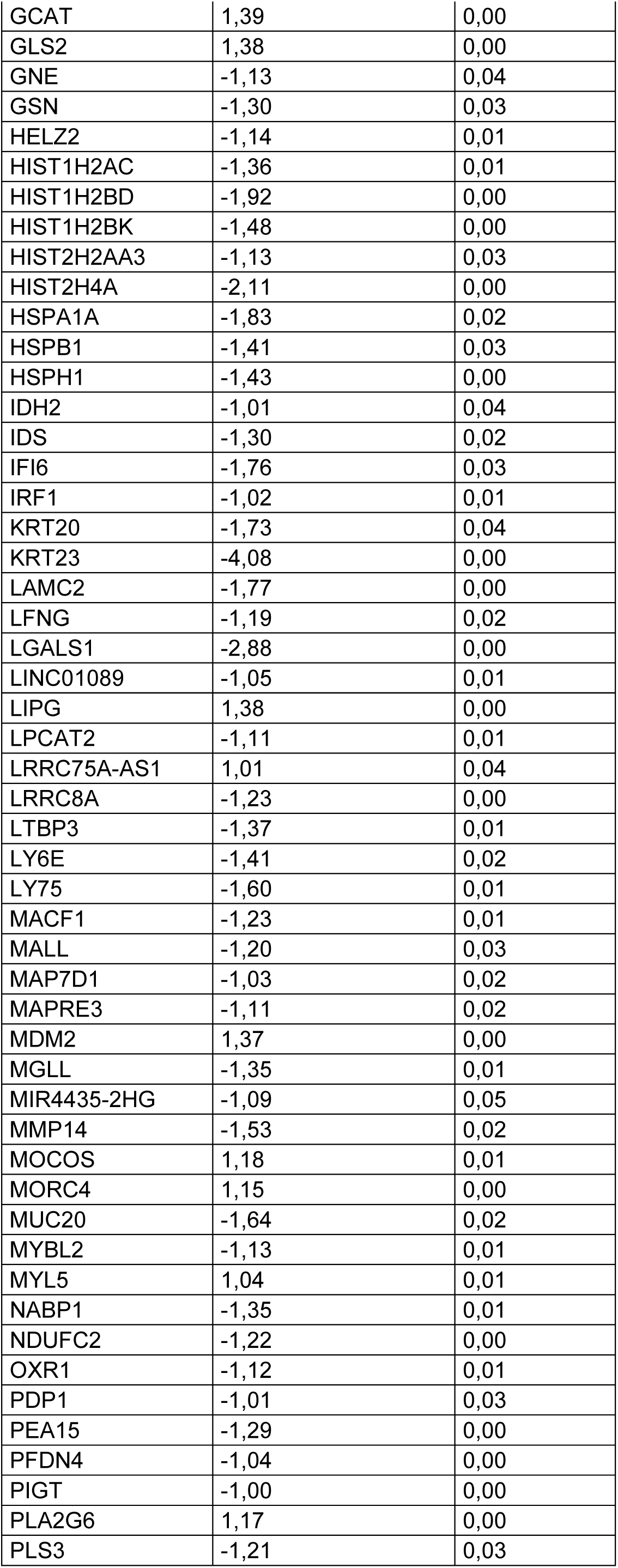
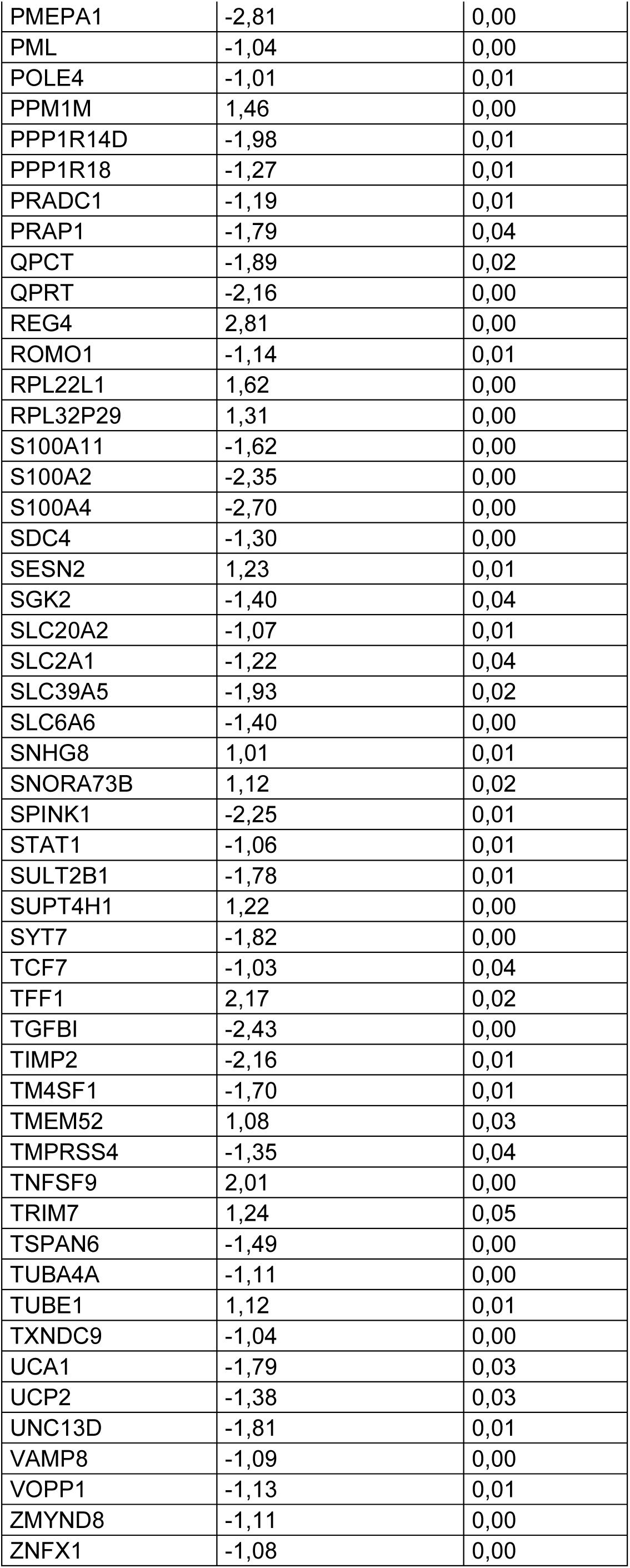
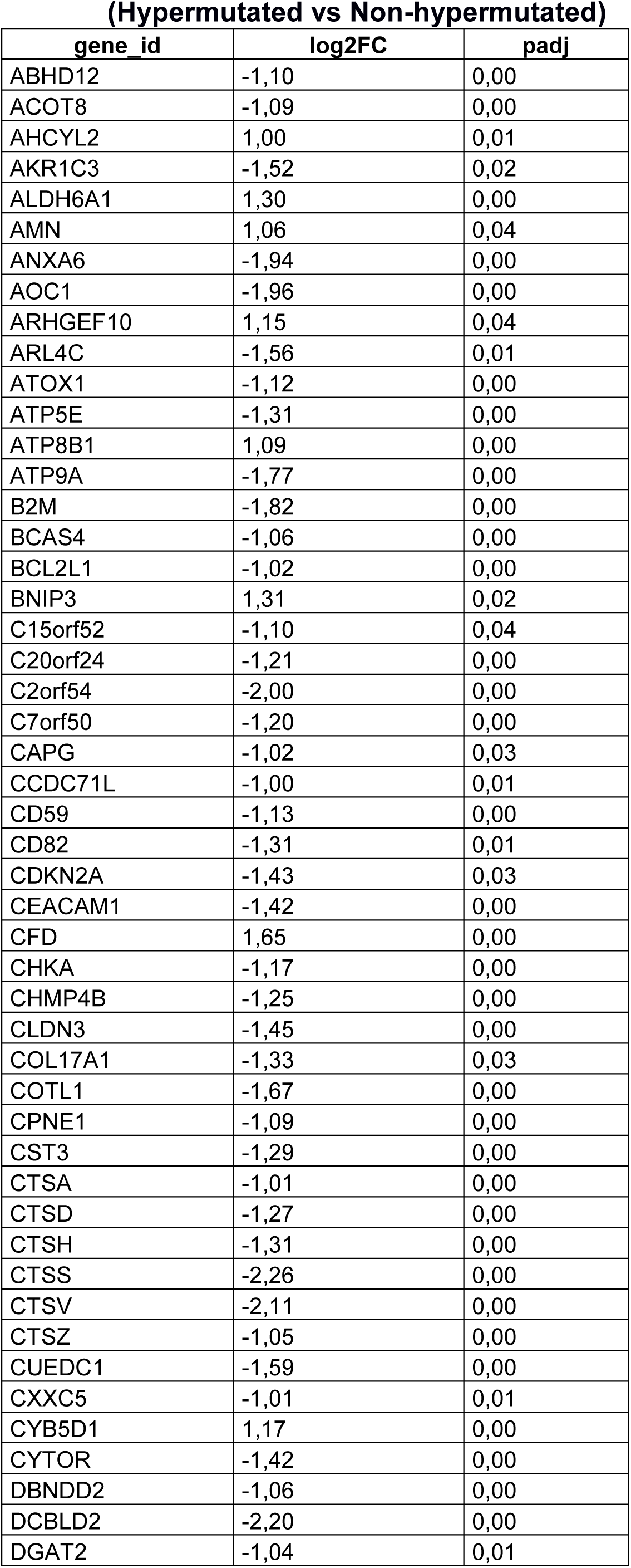
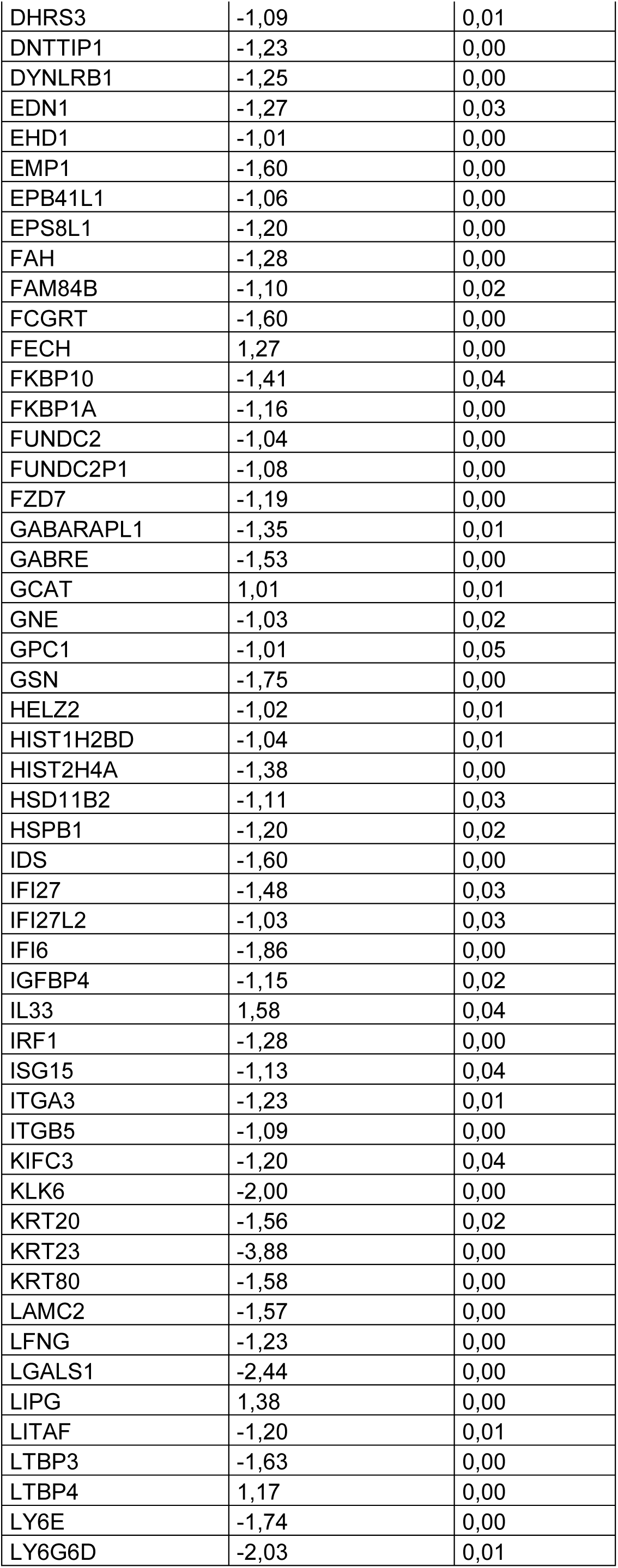
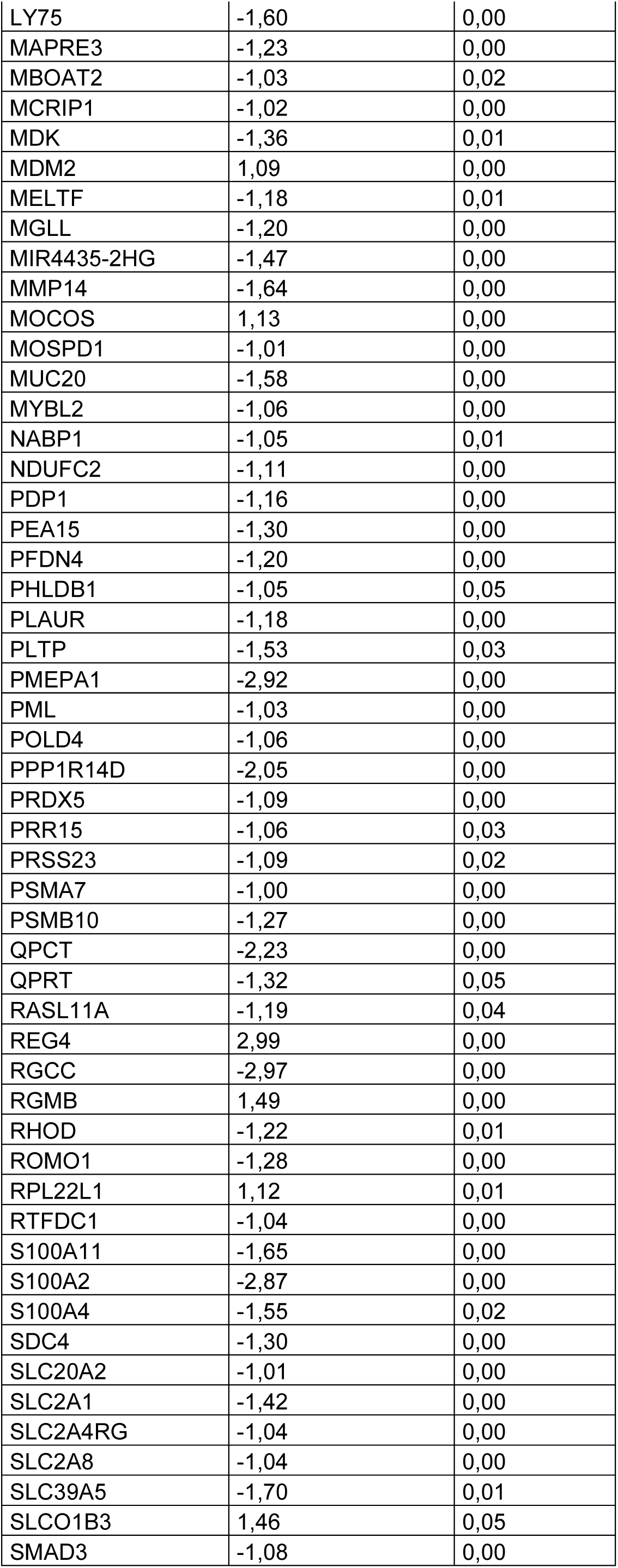
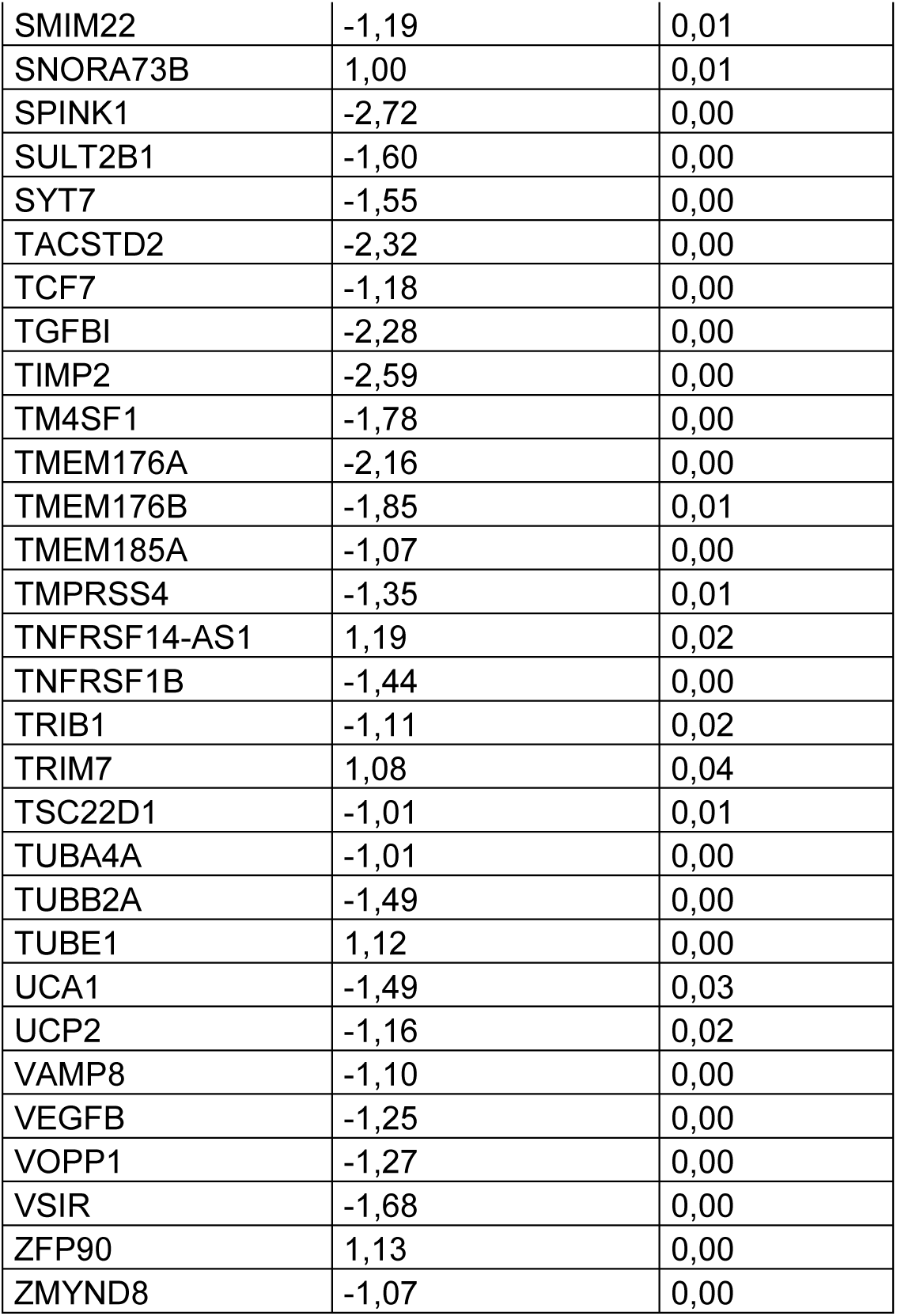
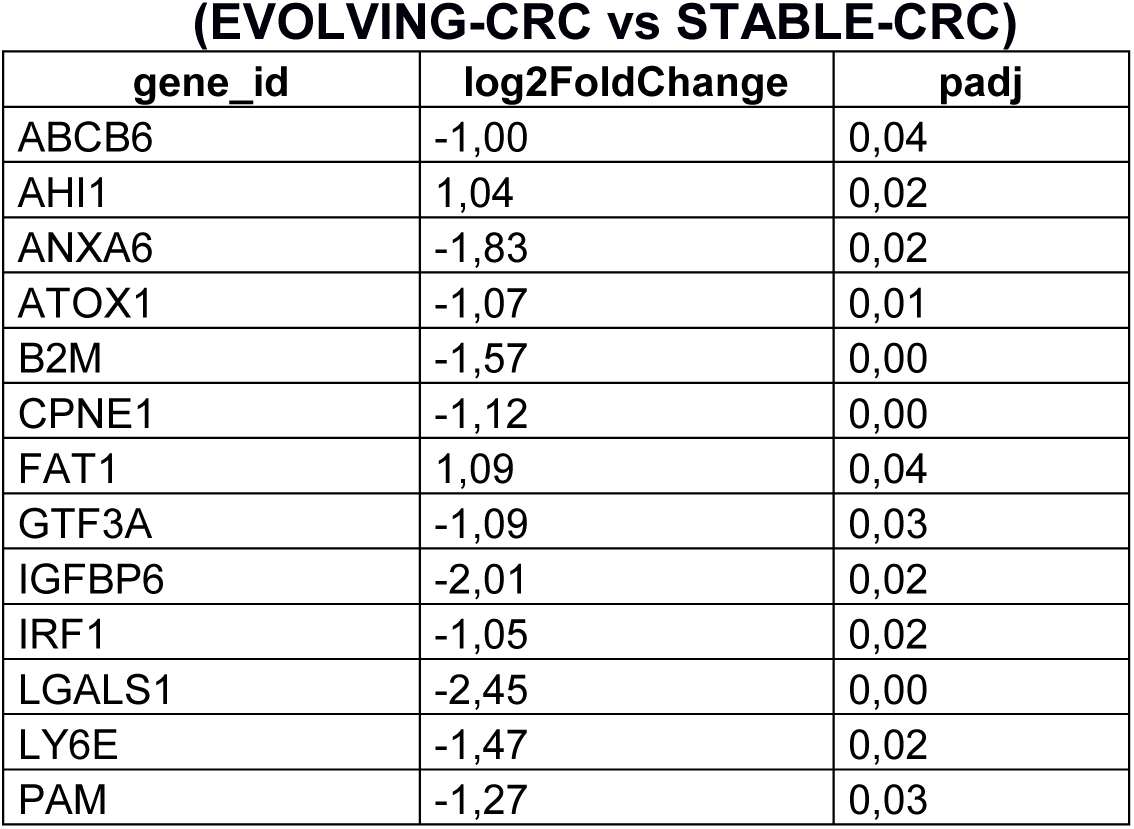
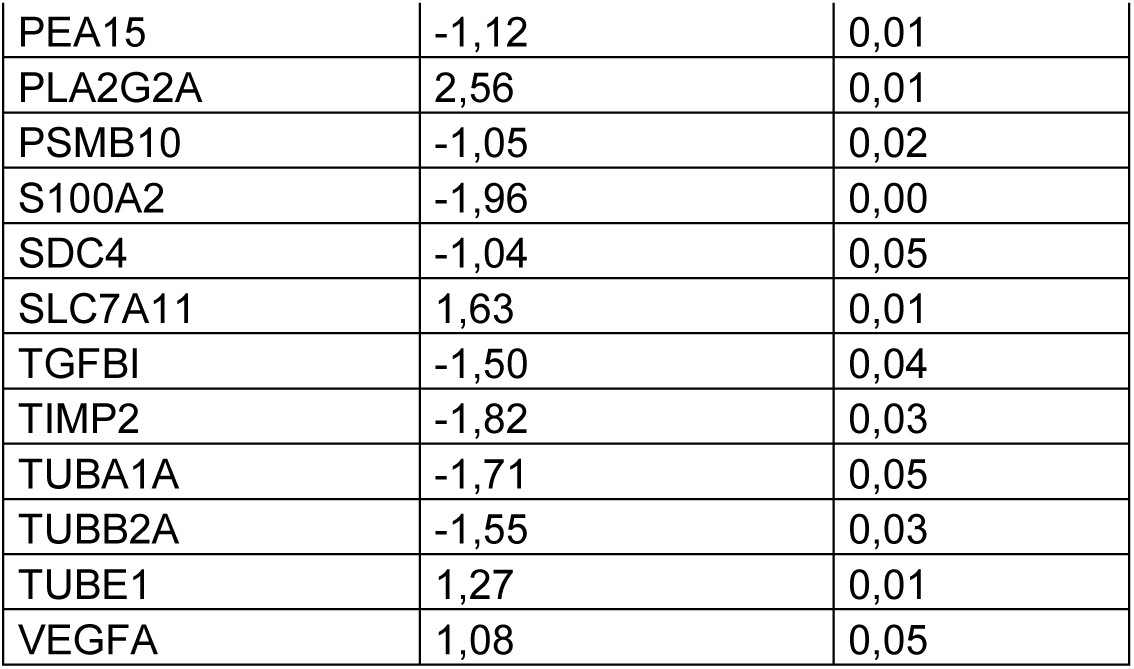
List of genes differentially expressed in the indicated cell lines.

**Fig. 8.**
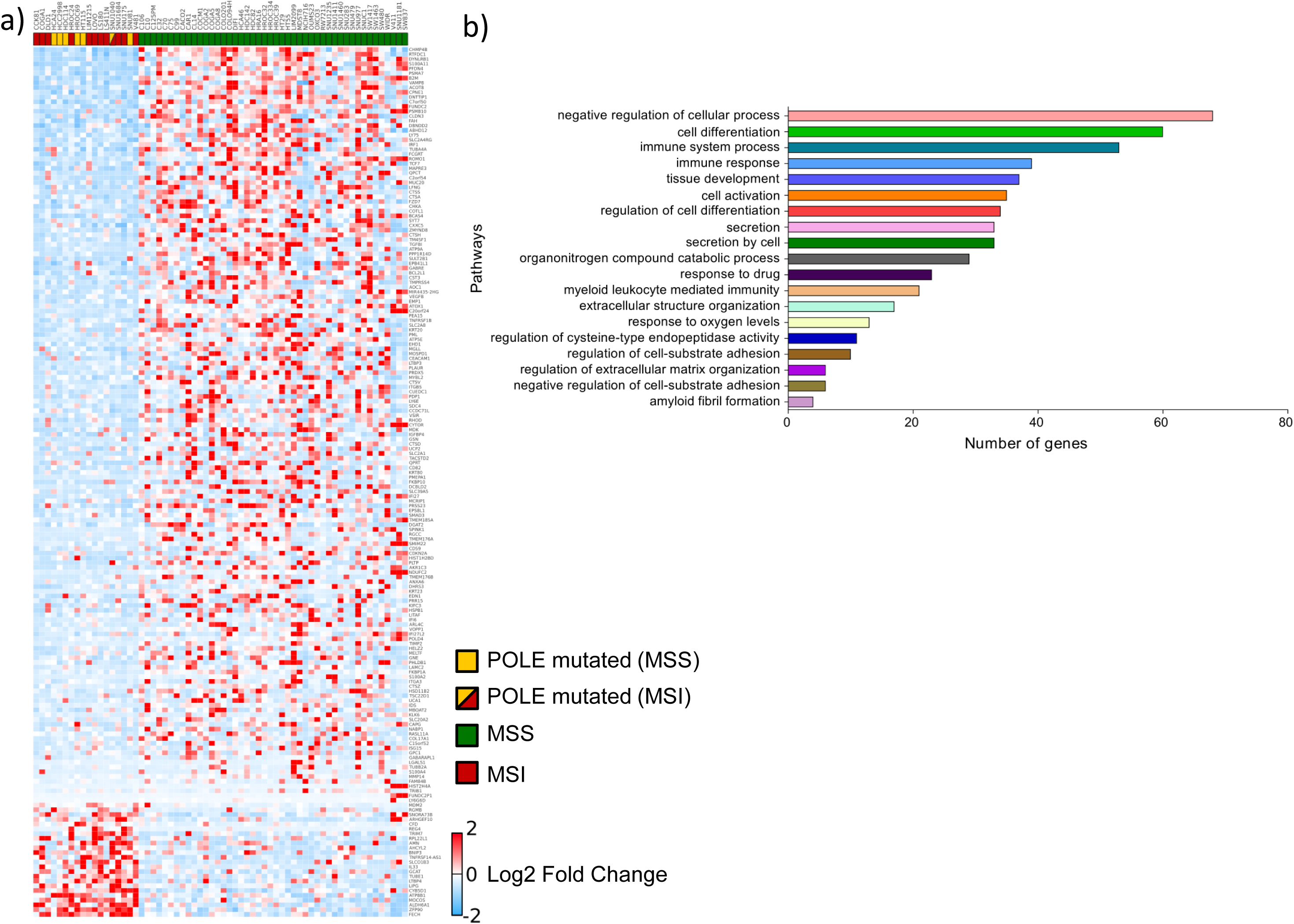
Transcriptional analysis of CRC cell lines. Differential expression analysis between hypermutated and non-hypermutated cells. a) 183 unique genes differentially expressed between hypermutated (MSI/*POLE*) versus non-hypermutated CRC cells (MSS). Log2 expression values, along with the mean change in expression are shown. b) Pathway analysis of genes differentially expressed between hypermutated versus non-hypermutated CRC cells using g:Profiler application (see Methods).

**Fig. 9.**
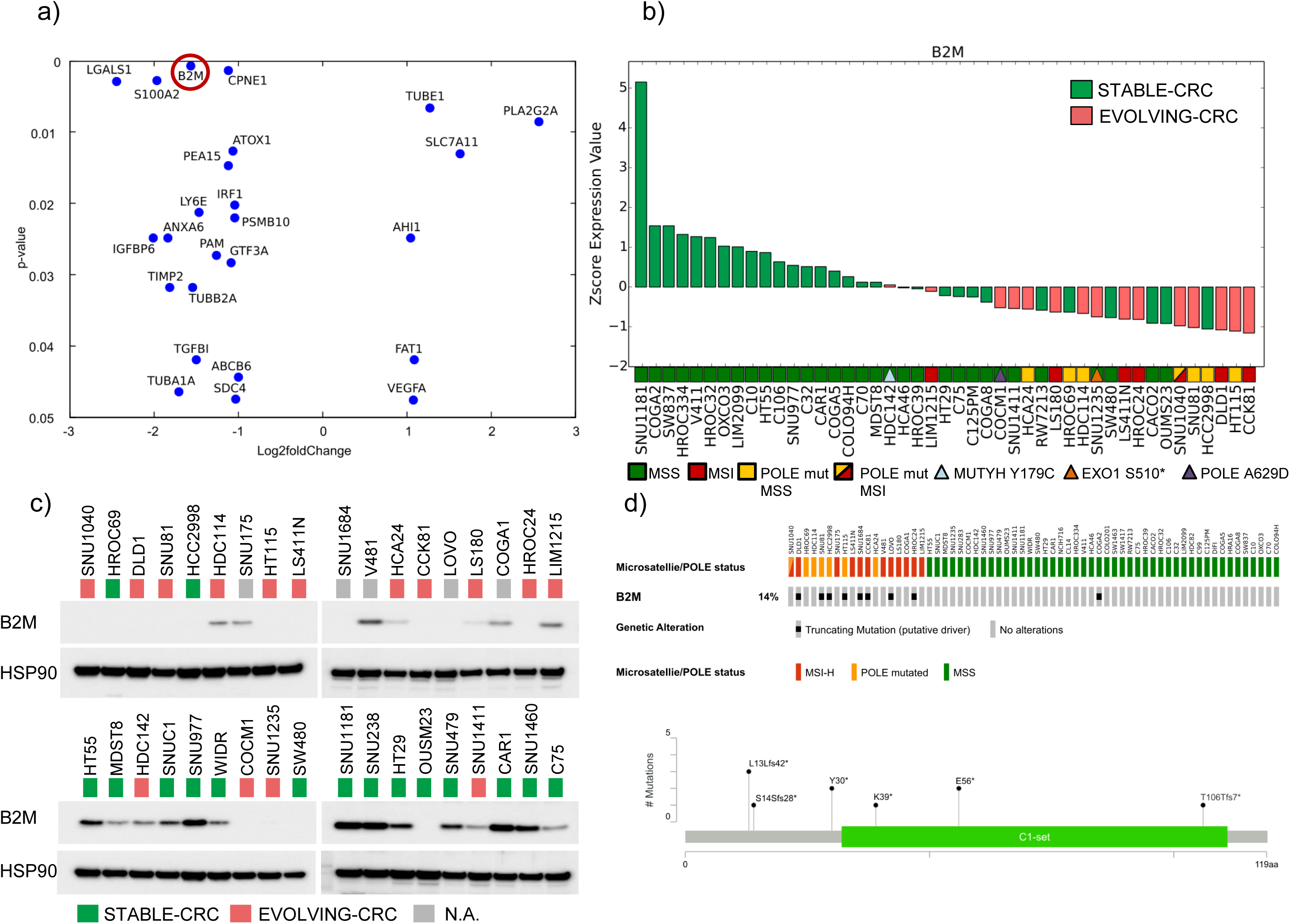
Beta2 microglobulin (B2M) expression is downregulated in EVOLVING-CRC. Transcriptional and protein levels of the B2M gene. a) Genes differentially expressed in EVOLVING-CRC relative to STABLE-CRC with a significant p-value (P < 0.05). b) Waterfall chart showing B2M expression at RNA level across a panel of 45 CRC cell lines. c) Western blot analysis of B2M expression. In gray are highlighted samples for which T90 sequencing were not available. Blots were reprobed with anti-HSP90 antibody to confirm equal loading. d) *B2M* gene alterations on 64 CRC cell lines at T0 (upper panel) and codon affected (lower panel).

**Table 5.**
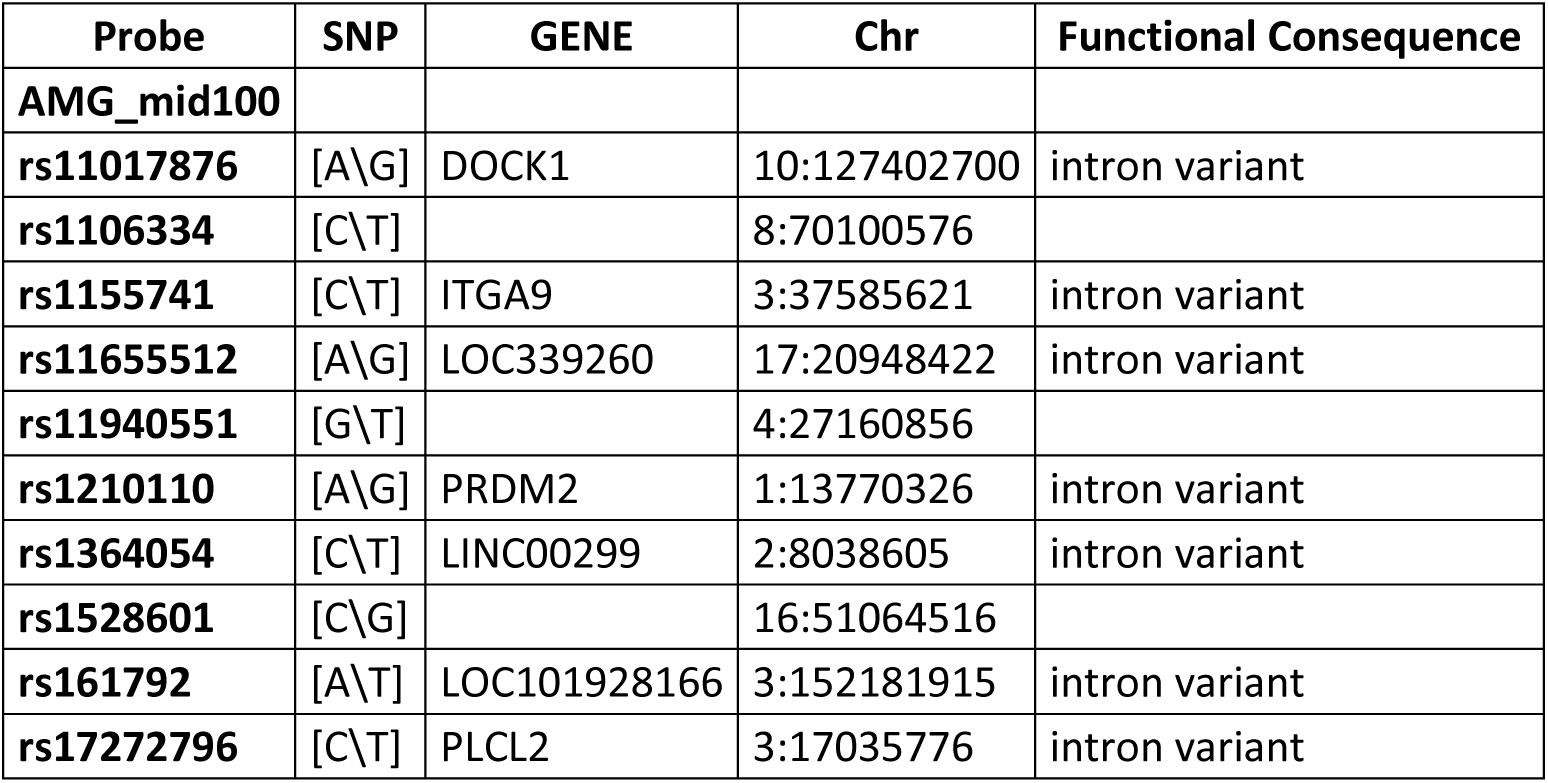
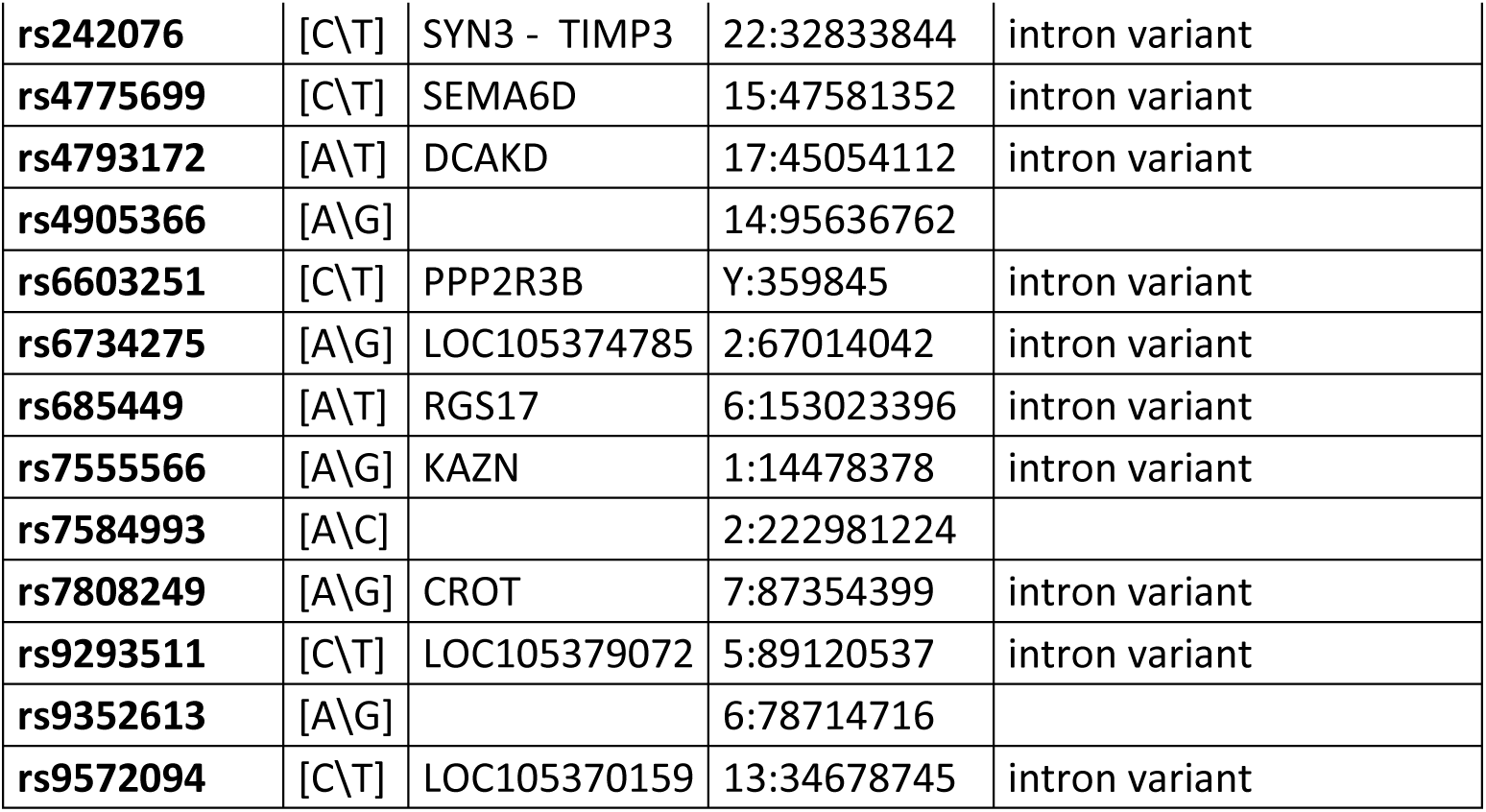
List of SNPs used to identify patient-derived xenografts.

## Discussion

In the past decade it has become clear that most human tumours are highly molecularly heterogeneous, and this affects prognosis and the emergence of therapeutic resistance (40). How tumour-specific somatic variations can lead to distinct neoantigen profiles and ultimately to immune surveillance has also been partially elucidated. The number of neoantigens depends on several factors. For example lung cancers associated with smoking habits have high levels of mutations (41) (42), whereas the development of skin melanomas is correlated with UV light mediated mutagenicity (43). Both smoking and UV exposure occur during defined periods and their mutagenicity is transient, leading to high -but relatively stable-mutational profiles (44) (45). Another class of tumours with high mutational burden is characterized not by exposure to external carcinogens, but rather by the intrinsic inability of tumour cells to efficiently repair DNA. The latter is due to epigenetic or genetic alterations in key effectors of DNA repair pathways, rather than acute or chronic carcinogen exposure. In this work, we used CRC as a model system to understand whether, and to what, extent alterations of DNA repair pathway components modulate neoantigen profiles over time *in vitro* and *in vivo*. Tumours carrying alterations affecting DNA repair genes maintained their molecular characteristics over time and, in most instances, the functional consequence of those alterations is continuous and propagated at every generation. An exception was represented by two *POLE* mutant CRC cell lines (HROC69 and HCC2998) which despite having high mutational burden did not appreciably evolve over time. The reason(s) for this phenotype are presently unclear. Interestingly these two *POLE* mutant cells that evolved poorly over time had less marked mutational signatures, possibly suggesting that, in these models, polymerase defects may undergo some form of functional compensation.

The longitudinal analysis of cell and PDX models highlighted several aspects. For example, MSI and POLE mutated tumours tended to acquire SNV or short insertions/deletions over time. These alterations can lead to novel putative neoantigens potentially trigger the host immune system. In addition to well-known DDR genes (*MLH1, MSH2, MSH6, PMS2, POLE*), our study, indicate that other genes involved in DNA repair pathway may lead to accumulations of mutations possibly translating in novel epitopes. *EXO1* and *MUTYH* are two of such examples. Profiling of these genes in the clinical setting may help to intercept tumours not classified as unstable or with hypermutator phenotype but nevertheless continuously evolving and accumulating mutations.

Our analysis suggests that in parallel to mutations gains, loss of variants also occurs during cell propagation. Our data indicate that in hypermutated CRCs, including MSI and *POLE* mutated models expanded *in vitro*, these events are mainly confined to subclones. A limitation of this study is that longitudinal characterization of lost and gained mutations *in vitro* could be influenced by sampling of cell populations during cell passaging. We also report that in the propagation of PDX, possibly due to selection imposed by the microenvironment, not only subclonal but also clonal variants emerge *de novo* over time. Based on these results, we speculate that in CRC patients with DNA repair defects metastatic seeding or therapeutic debulking can lead to the emergence of new subsets of clonal neoantigens. This could have implications for the development of therapies relying on the presence of clonal neo antigens, such as ICP, CAR-T and vaccines.

Both cell lines and PDX have been widely employed to test anticancer compounds (46-48), however experimental reproducibility has occasionally been questioned (49, 50). The molecular evolvability that we find to occur during serial passaging of cells and PDX may partly account for the discrepant results obtained with these models (51-53).

A limitation of the present study is that it examined the evolution of cell lines and xenografts but cannot address the impact of the immune system in the evolutionary dynamics due to intrinsic limitations of the models we used.

Our data indicate that alterations in DNA repair genes facilitate acquisition of neoantigens. These novel putative epitopes can be recognized by the immune system. Accordingly, we confirm that CRCs with high number of mutations (hypermutated CRCs) selectively downregulate components of the neoantigen presentation process, such as *B2M*, thus restricting the ability of the host immune system to detect them. Our results further suggest that non hypermutated CRCs, that display fast evolving mutational and antigen profiles, also show downregulation of components implicated in neoantigen presentation. The differences in expression of molecules involved in immune functions we observed in the CRC models could have originated from adaption previously experienced in the patient as a mechanism of escape from negative pressure of the immune system related to the elevated neoantigens’ production rate.

## Conclusions

In summary, we identified, and functionally highlighted CRC subsets characterized by slow and fast genome evolvability. CRCs carrying alterations in genes involved in DNA repair (including *MLH1*, *MSH2*, *MSH6*, *MUTYH*, *EXO1*, and *POLE*) display dynamic neoantigen patterns that fluctuate over time. Furthermore, we find that in CRC cells and patient-derived tumour xenografts, DNA repair defects leading to high mutational burden and neoantigen evolvability are associated with inactivation or downregulation of antigen-presentation functions. Longitudinal monitoring the neoantigen landscape of CRC and other tumour types may have clinical implications. While tracking time-dependent neoantigen evolution in the tissue of cancer patients might be difficult or impossible to achieve, monitoring predicted neoantigens in circulating tumour DNA is already within reach. Accordingly, longitudinal liquid biopsies could be deployed to assess whether and how time and/or therapeutic regimens affect the mutational burden and the neoantigen profiles in individual patients. Neoantigen clonality profiles could be valuable to develop specific vaccines and deploy immunomodulatory molecules in the context of precision oncology.

## Declarations

### Ethics approval and consent to participate

All animal procedures were approved by the Ethical Commission of Candiolo Cancer Institute IRCCS, of the University of Turin and by the Italian Ministry of Health. They were performed in accordance with institutional guidelines and international law and policies. The number of mice included in the experiments, the inclusion/exclusion criteria and the observed tumour size limits were based on institutional guidelines. Guidelines limited us to using 6–8-week-old female and male NOD–SCID mice. Mice were obtained from Charles River. Informed consent for research use was obtained from all patients at the enrolling institution prior to tissue banking and study approval was obtained from the different centres.

### Consent for publication

Not applicable

### Availability of data and material

The datasets used in the current study are available from the corresponding author upon reasonable request.

## Authors’ contributions

A.B., G.R., A.L., and N.A. conceived and designed the study. A.L., N.A., A.M., C.C., V.A., M.M., A.Bart., and G.G. performed the experiments. G.R., G.C., C.N., L.N., and C.I. performed bioinformatics analysis. A.B., G.R., A.L., and G.G. interpreted the data. L.T. and A.Ber. provided samples and interpreted the data. A.B., G.R., and A.L. wrote the manuscript with input from all authors. All authors read and approved the final manuscript.

## Acknowledgements

We acknowledge Francesco Galimi for providing gDNA and RNA from PDXs. We thank Daniela Cantarella for performing RNAseq. We thank all members of our laboratory for valuable suggestions, critical review of the data, and the manuscript, as well as all laboratory technicians for their help.

---

**Supplementary Fig. 1.**
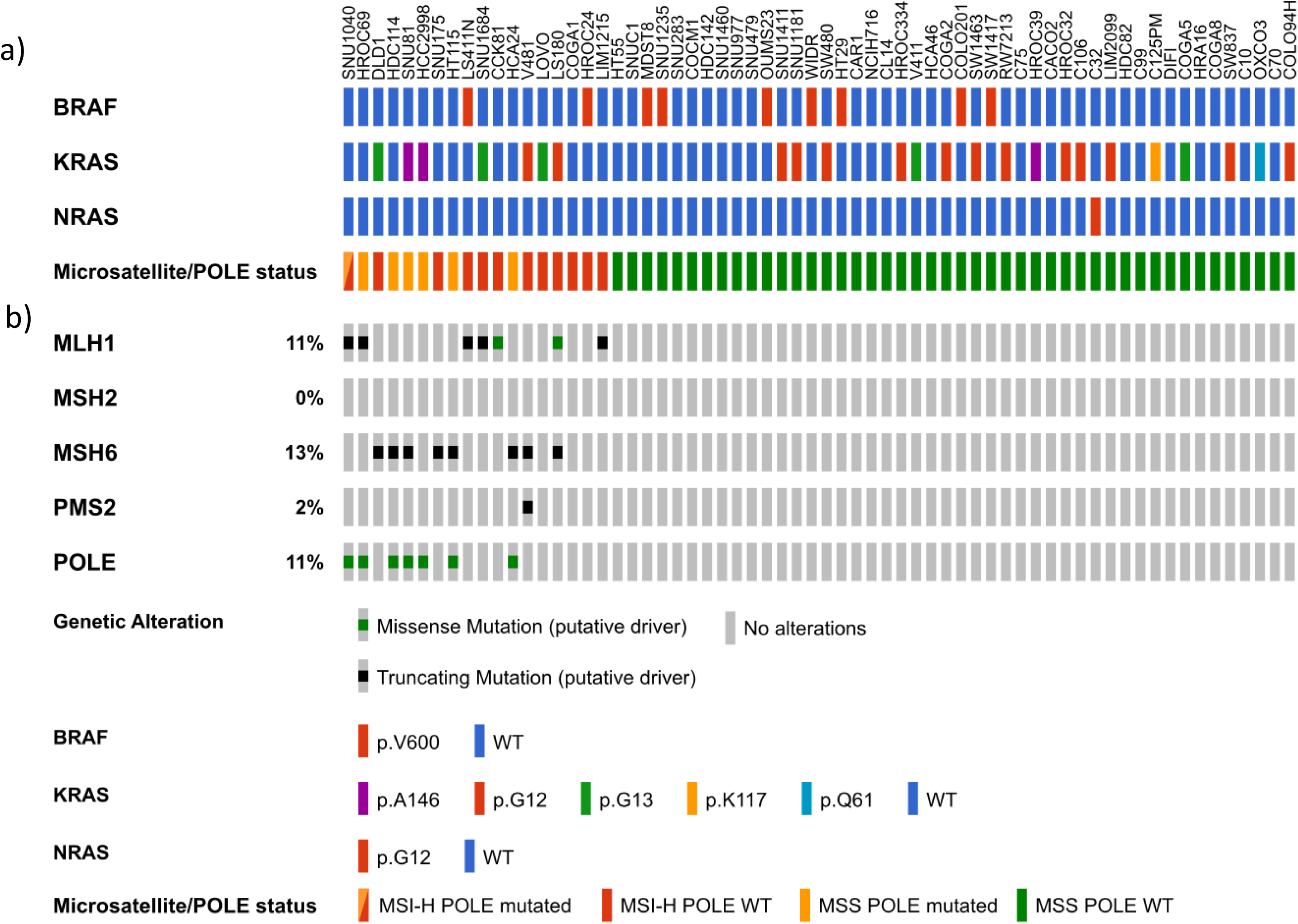
Genomic features of 64 CRC cell lines. Molecular characterization of the indicated CRC models at T0 using cbioportal oncoprint graphic representation. a) Schematic diagram showing *BRAF*, *KRAS*, *NRAS* and Microsatellite/*POLE* status of CRC cell lines. b) Schematic diagram showing genetic alterations in MMR genes and DNA proofreading polymerase *POLE* in CRC cell lines.

**Supplementary Fig. 2.**
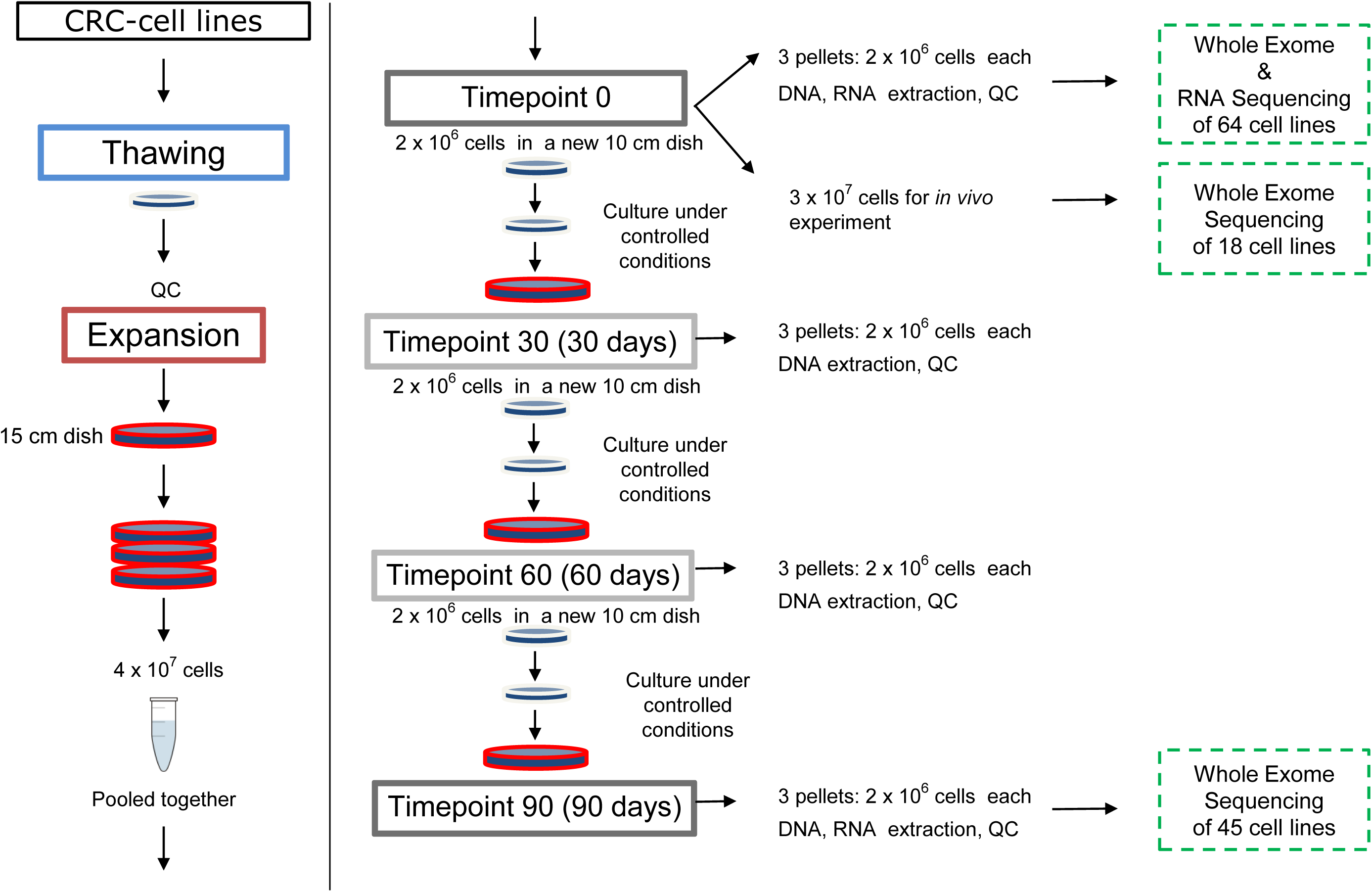
Outline of the experimental workflow to assess evolution of CRC cell lines *in vitro* and *in vivo*. CRC cell lines were thawed and kept in culture for 90 days. WES and RNAseq were performed at the beginning of the experiment (T0) for 64 cell lines. Forty-five samples collected at T90 were also subjected to WES. In a few instances at T0 an equal number of cells was injected in two immunodeficient mice. When tumours reached approximately a volume of 1000m^3^ in size, they were excised and sequenced.

**Supplementary Fig. 3.**
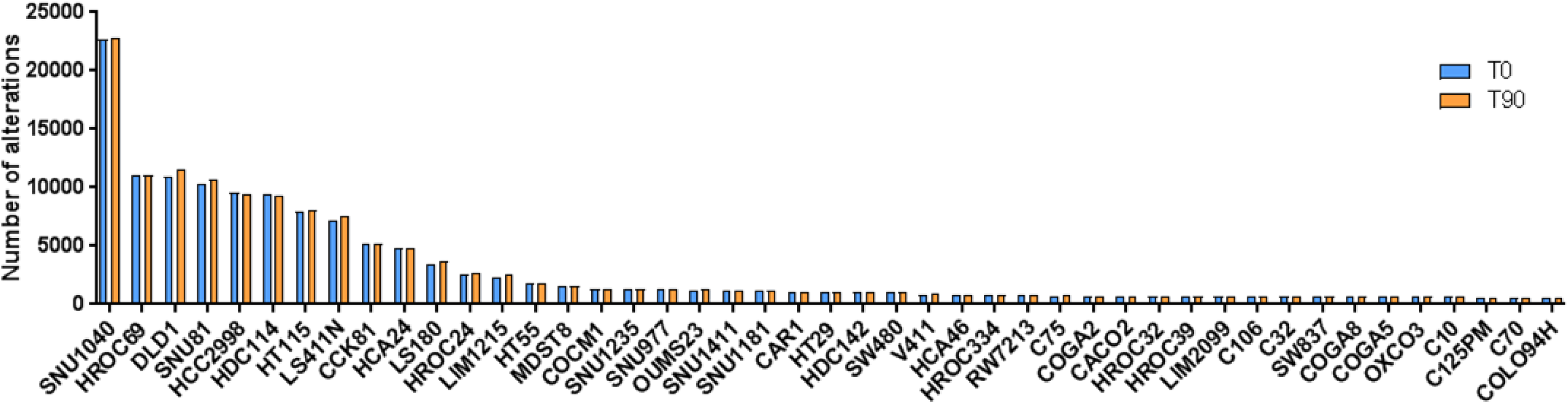
Genetic alterations in CRC cell lines at T0 and T90. Bar chart showing the number of alterations at the beginning of experiment (T0) and after 90 days of culture (T90). All the variants are called against the hg38 reference.

**Supplementary Fig. 4.**
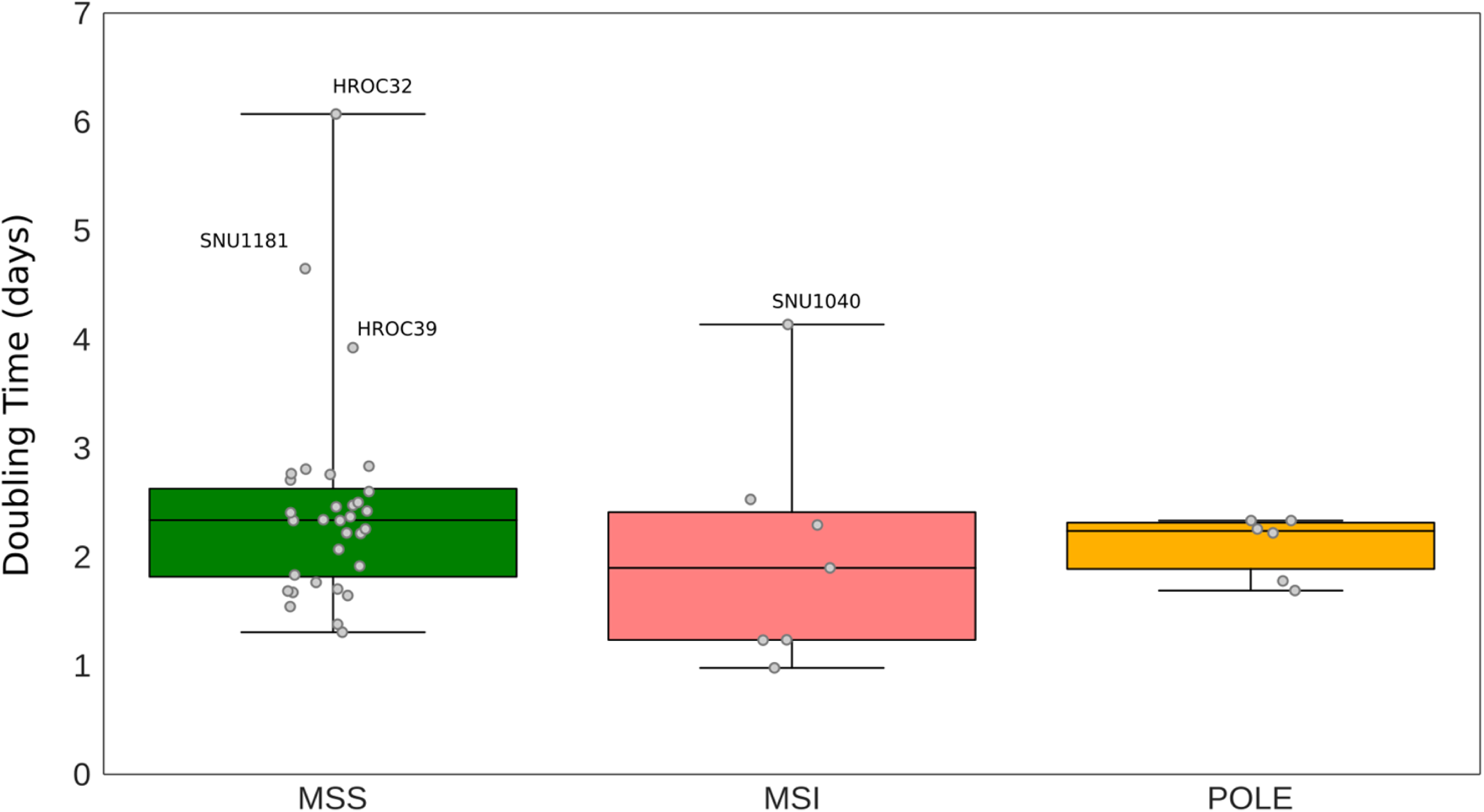
Doubling time of MSS, MSI and POLE mutated cell lines. Cell doubling time in MSS, MSI and MSS *POLE* mutant cells. The number of days per group is shown. The centre line of each box plot indicates the median.

**Supplementary Fig. 5.**
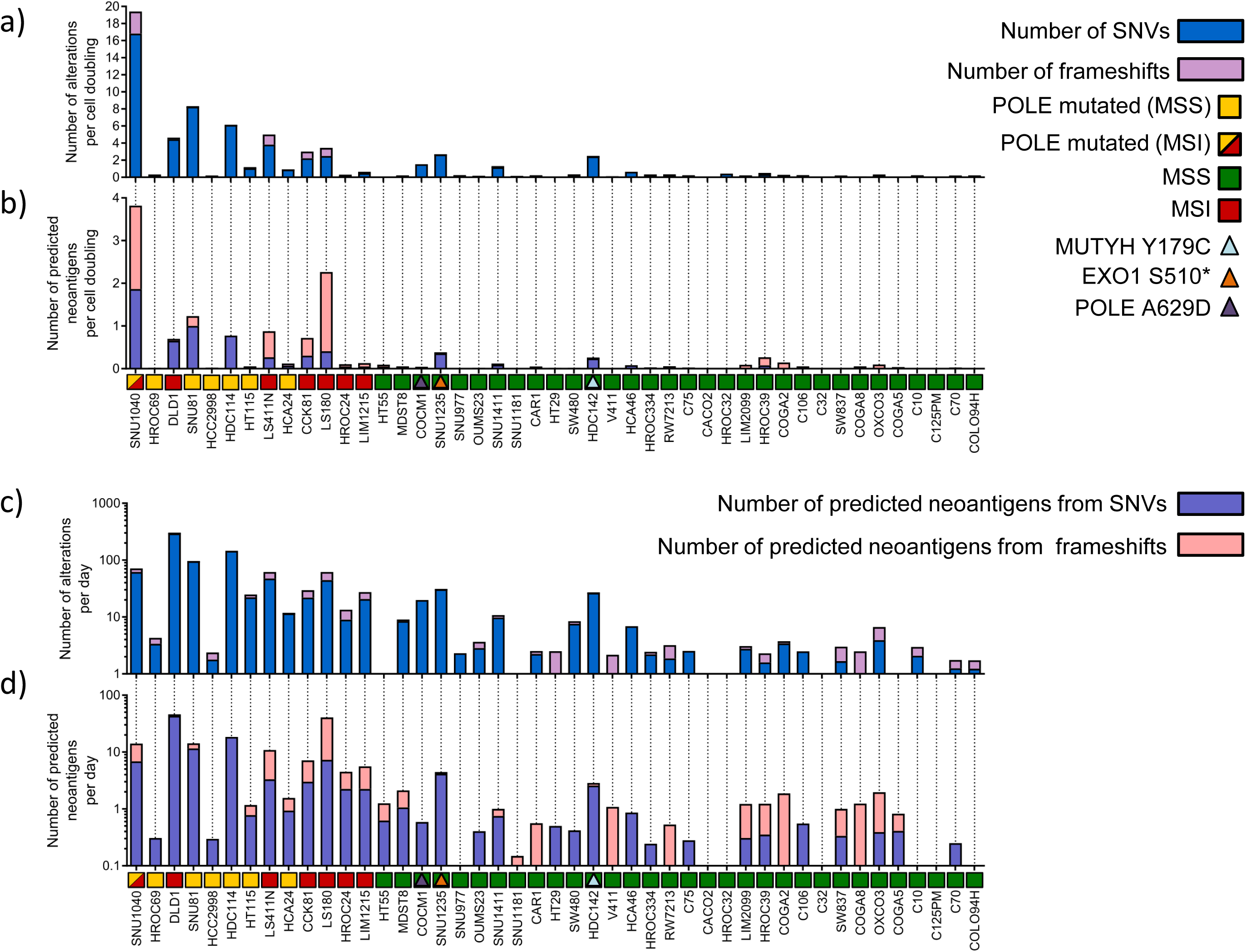
*In vitro* evolution of mutational landscape in CRC cell line normalized to the doubling time. Mutational characterization of CRC cells after 90 days of culture (T90) normalized to the doubling time. a) The bar chart shows the number of new alterations acquired at T90 (absent at T0) normalized to the cell doubling time. b) The number of predicted neoantigens per cell doubling (see Methods) is shown. Each bar represents putative neoepitopes derived from SNVs and frameshifts. c) The bar chart shows the number of new alterations per day acquired at T90 (absent at T0). d) The number of predicted neoantigens acquired per day (see Methods) is listed. Each bar represents putative neoepitopes derived from SNVs and frameshifts.

**Supplementary Fig. 6.**
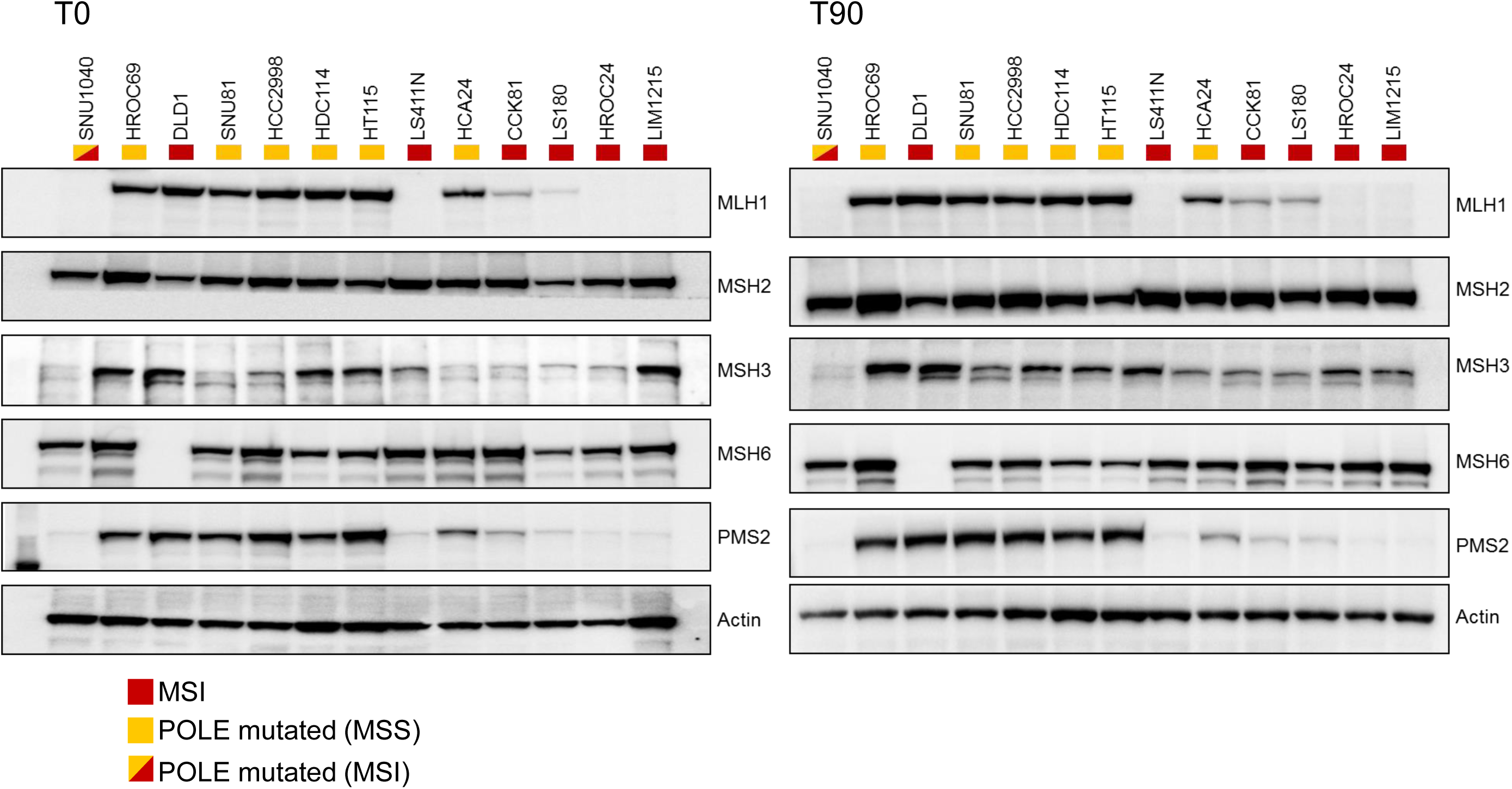
MMR proteins expression in cell models. Western blot analysis on hypermutated samples (MSI and/or POLE mutant cell lines) to assess the status of the MMR proteins at Time 0 (T0) and Time 90 (T90).

**Supplementary Fig. 7.**
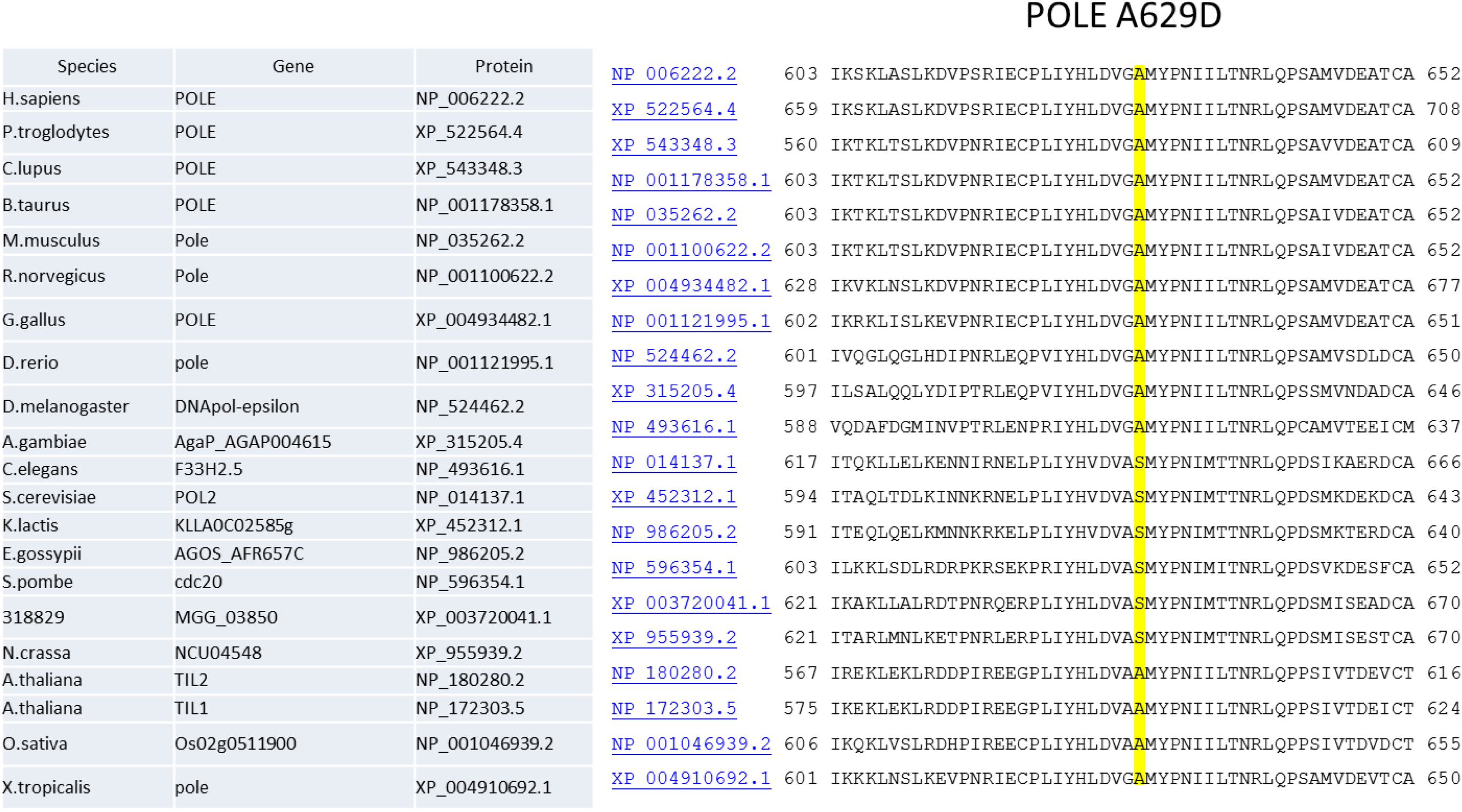
POLE protein sequence alignment across species. The yellow box indicates alanine (A) 629, which is highly conserved across species but is mutated to aspartic acid (D) in the COCM1 cell line.

**Supplementary Fig. 8.**
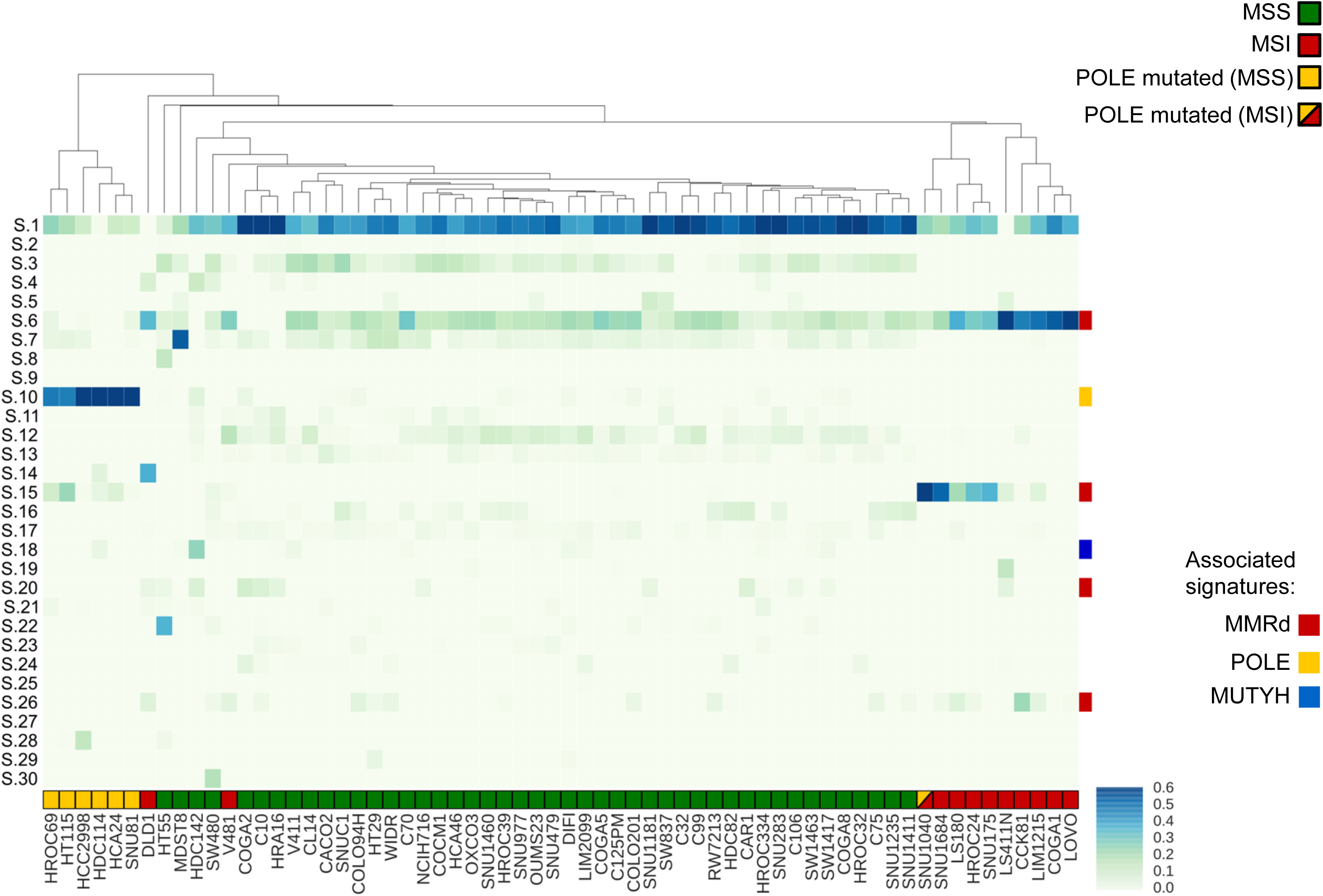
Analysis of mutational signatures in CRC cell lines. Heatmap and clustering of 30 cancer associated signatures across 64 CRC cell lines at T0. Signatures associated with MMR-deficiency (6, 15, 20 and 26), POLE mutations (10) and MUTYH-associated polyposis (18) are highlighted. Analysis and clustering were performed as reported in Methods.

**Supplementary Fig. 9.**
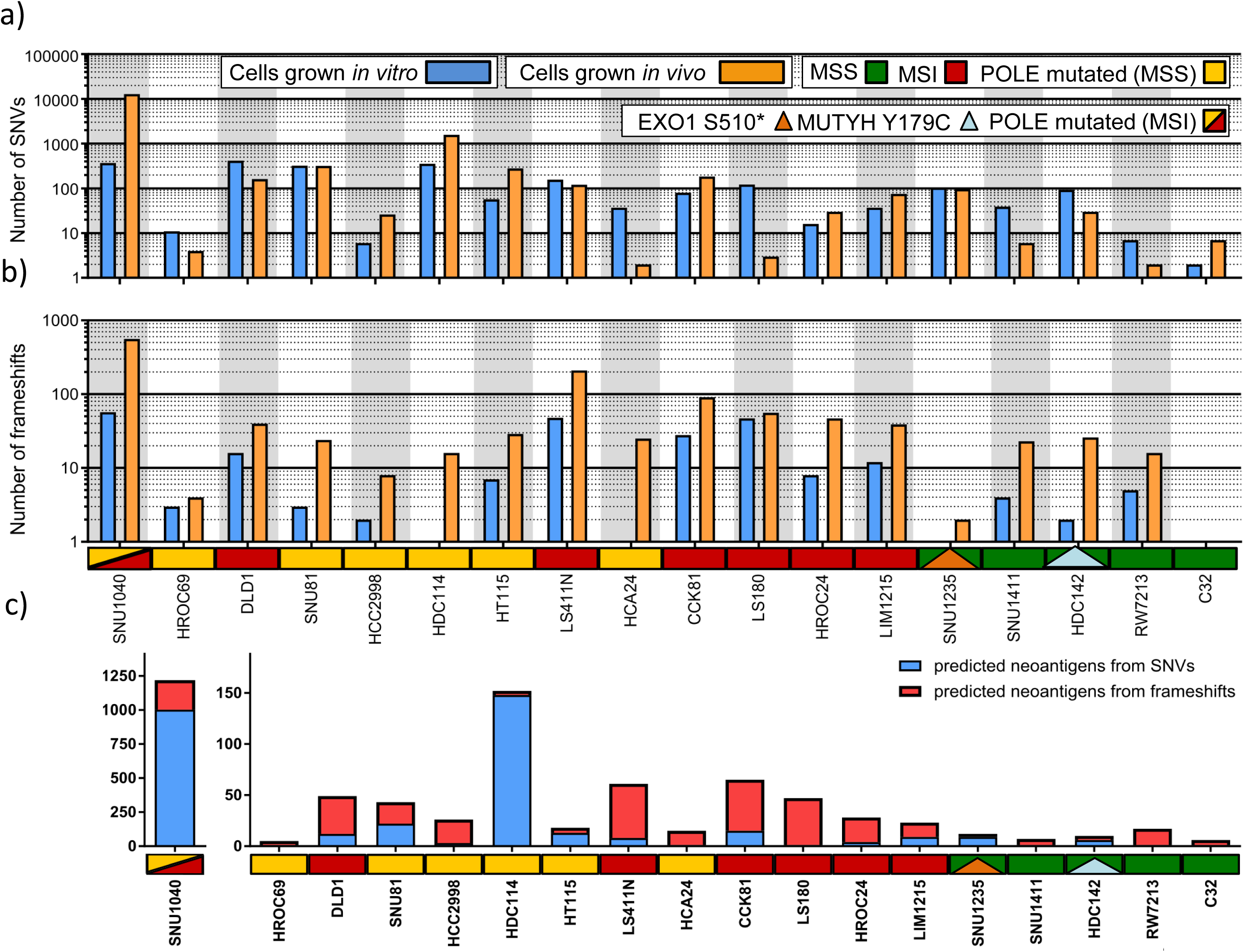
Comparison of mutational profiles in cell lines grown *in vitro* and *in vivo*. Alterations acquired by the indicated cell lines after 90 days in cell culture and upon transplantation in mice. a) Number of SNVs/Mb acquired *in vitro* (blue) and *in vivo* (yellow). b) Number of frameshifts acquired *in vitro* (blue) and *in vivo* (yellow) c) Number of predicted neoantigens in CRC cell lines after injection in mice (See Method for details).

**Supplementary Fig 10.**
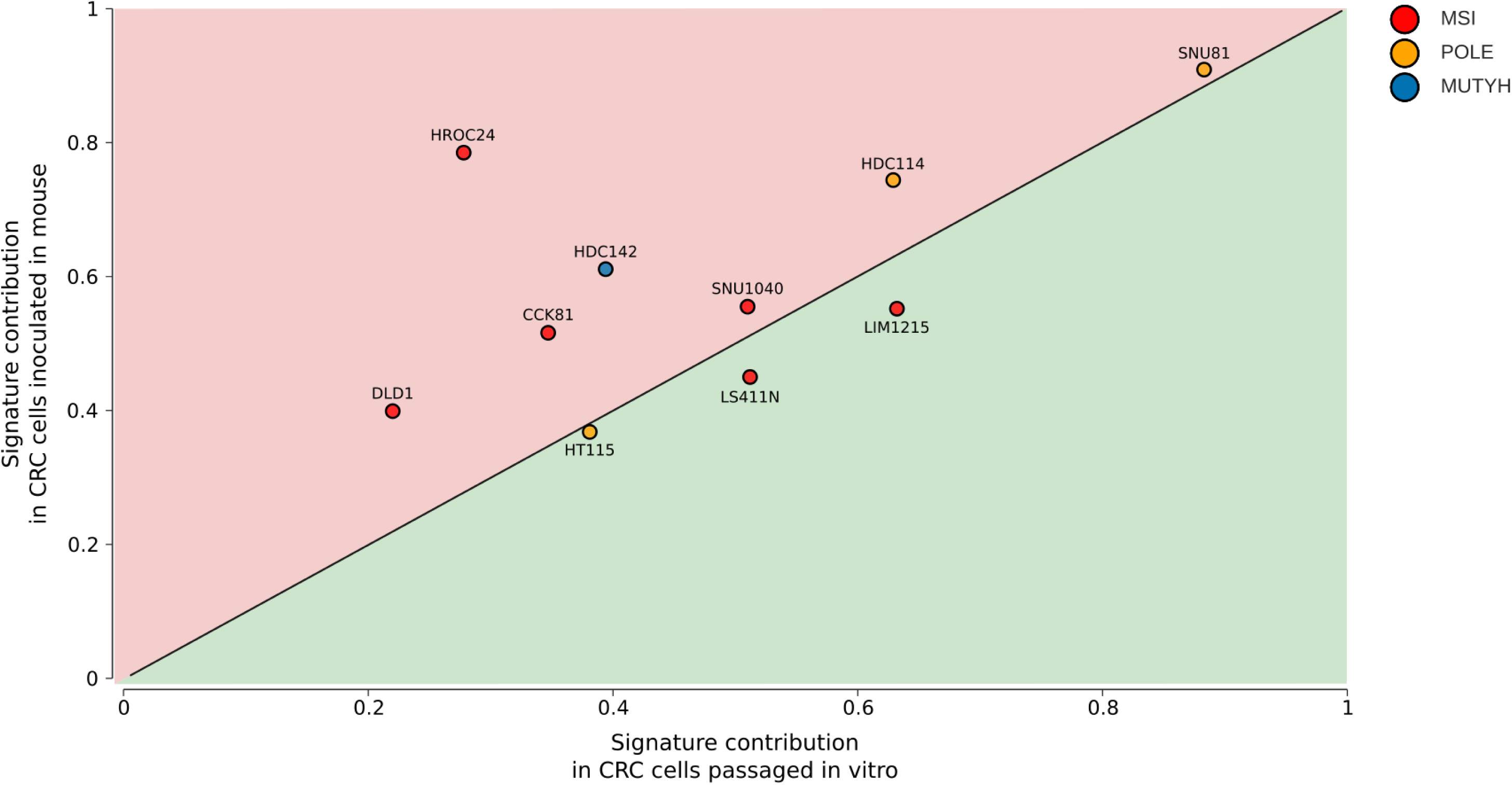
Comparison of signatures contribution between CRCs models propagated *in vitro* and *in vivo*. The contributions of the associated signatures based on the molecular status detected *in vitro* and *in vivo* were compared in each model. For the MSI samples we compared the maximum contribution provided by signatures 6, 15, 20 or 26 (linked to MMRd). For the MSS POLE mutant cells, contribution was from signature number 10. For the cell line carrying MUTYH biallelic alterations contribution was from signature 18.

**Supplementary Fig. 11.**
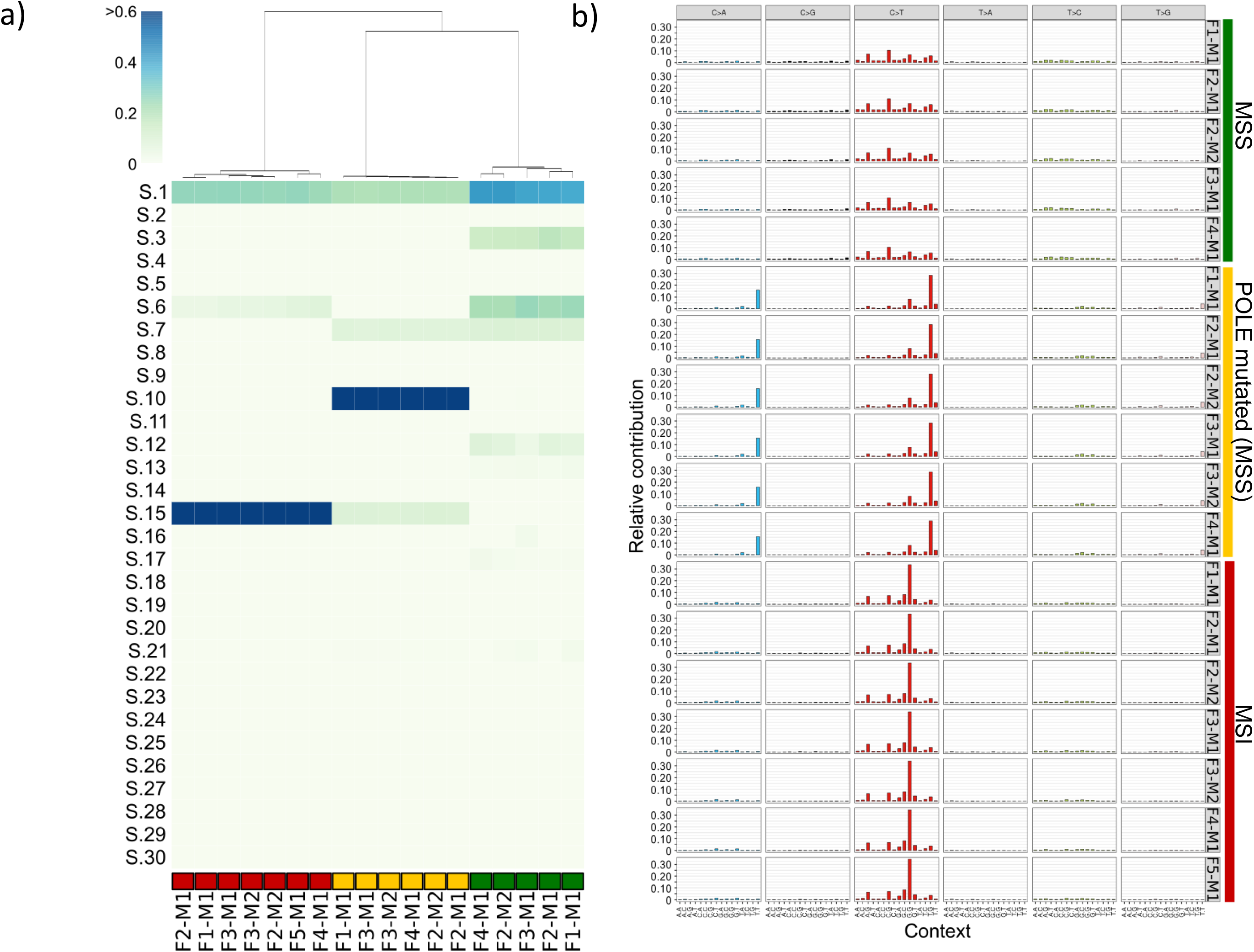
Analysis of mutational signatures in patient-derived xenograft. Contributions of 30 cancer associated signatures in patient-derived CRC xenografts. (a) Clustered heatmap according to signatures contribution in the indicated PDXs. (b) Signature profiles in PDXs using the six substitution subtypes: C>A, C>G, C>T, T>A, T>C, and T>G. Analysis and clustering were performed as reported in Methods.

**Supplementary Fig. 12.**
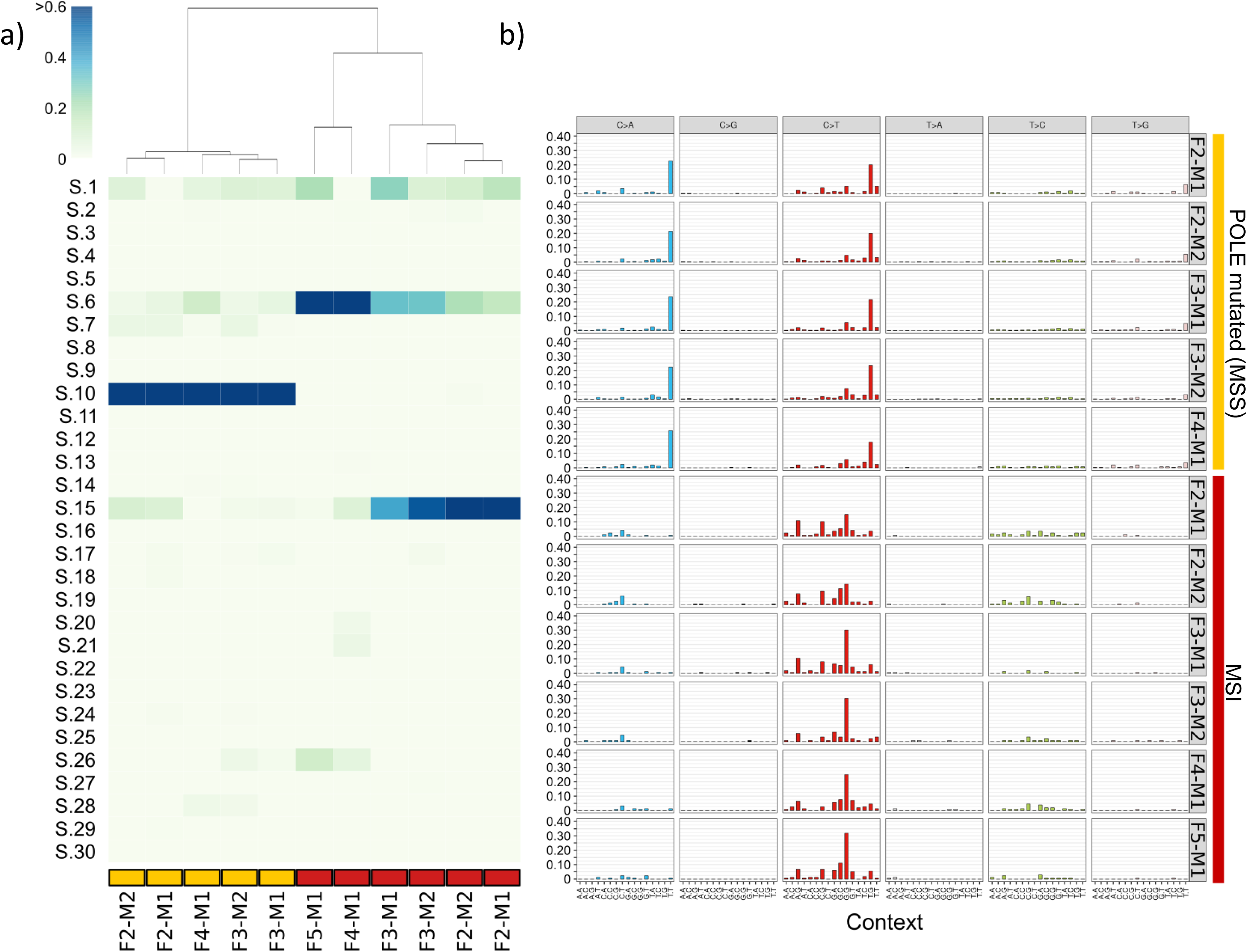
Mutational signatures acquired during propagation of patient-derived xenografts. Contributions of 30 cancer-associated signatures and signature profiles across evolving PDXs. Genomic variants of the individual PDXs were compared to the corresponding previous generation to infer signatures contributions. a) Clustered heatmap of signature contributions during evolution of the indicated PDX generations. b) Signature profiles acquired in each PDX generation using the six substitution subtypes: C>A, C>G, C>T, T>A, T>C, and T>G. Alterations of each sample were inferred comparing two consecutive generations (see Method for detailed information).

**Supplementary Fig. 13.**
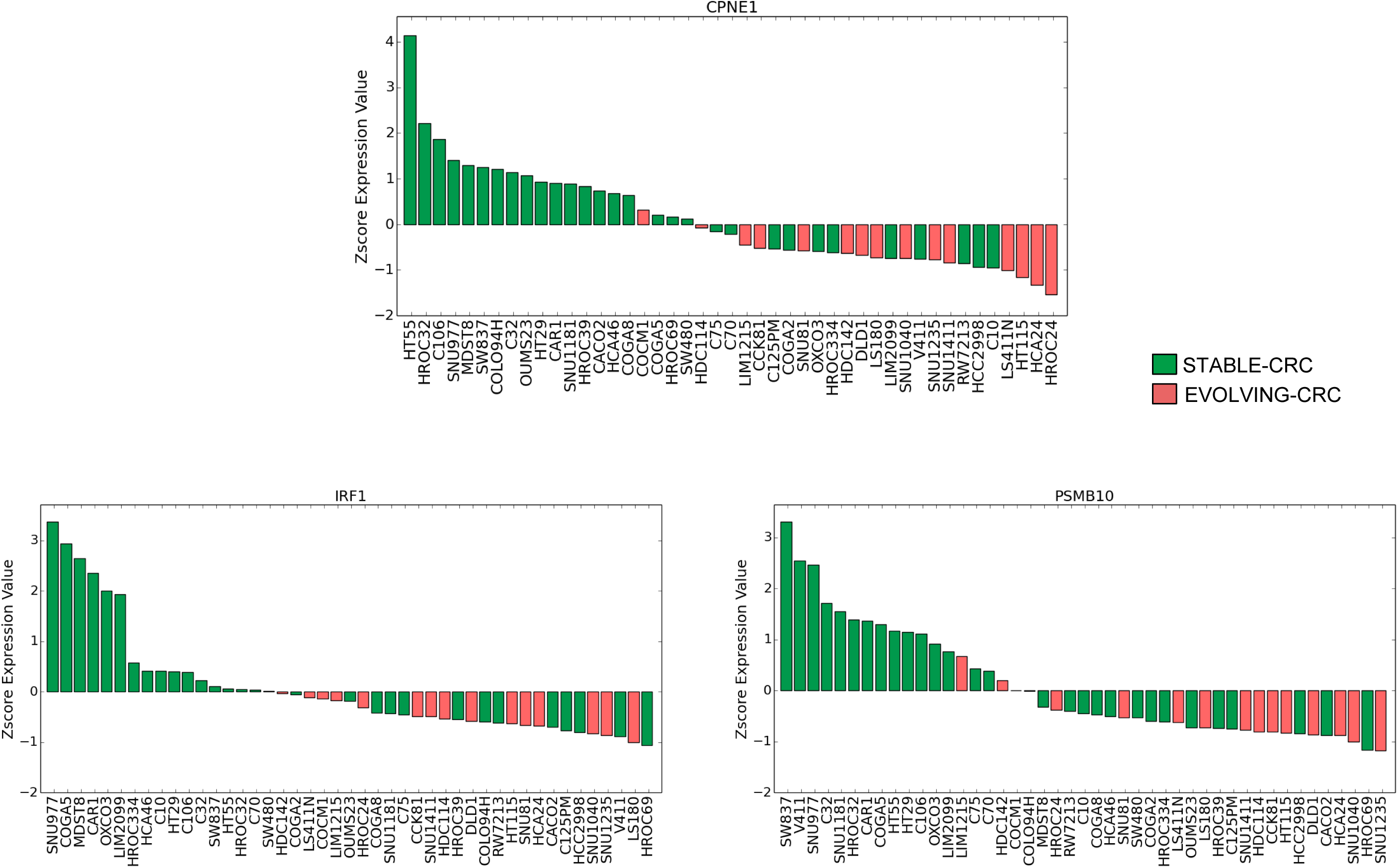
Gene differentially expressed in EVOLVING-CRC. Waterfall charts show the z-score expression values of IRF1, CPNE1 and PSMB10 across a panel of 45 CRC cell lines.

**Supplementary Fig. 14.**
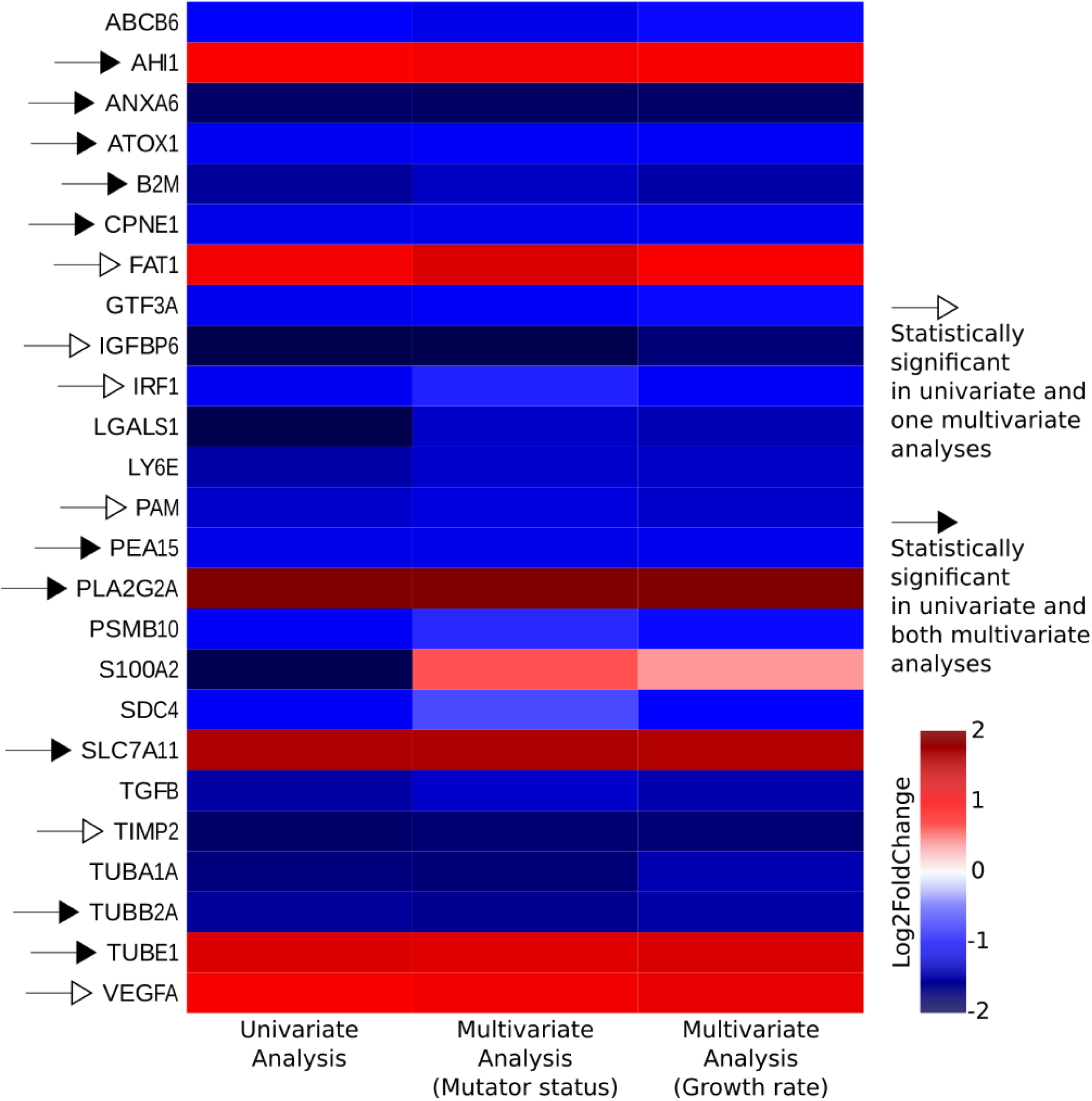
Gene differentially expressed in EVOLVING-CRC in univariate and multivariate analyses. Log2 fold-change of the genes listed in Fig 9a according to univariate and multivariate analyses considering the mutator status or the growth rates of the cells.

